# Transcriptomic signatures of ageing vary in solitary and social forms of an orchid bee

**DOI:** 10.1101/2020.07.30.228304

**Authors:** Alice C. Séguret, Eckart Stolle, Fernando A. Fleites-Ayil, José Javier G. Quezada-Euán, Klaus Hartfelder, Karen Meusemann, Mark Harrison, Antonella Soro, Robert J. Paxton

**Author notes:** Correspondence: A. C. Séguret (,), Robert J. Paxton.

## Abstract

Eusocial insect queens are remarkable in their ability to maximise both fecundity and longevity, thus escaping the typical trade-off between these two traits. In species exhibiting complex eusocial behaviour, several mechanisms have been proposed to underlie the remoulding of the trade-off, such as reshaping of the juvenile hormone pathway, or caste-specific susceptibility to oxidative stress. However, it remains a challenge to disentangle the molecular mechanisms underlying the remoulding of the trade-off in eusocial insects from caste-specific physiological attributes that have subsequently arisen due to their different life histories. Socially plastic species such as the orchid bee *Euglossa viridissima* represent excellent models to address the role of sociality *per se* in longevity as they allow direct comparisons of solitary and social individuals within a common genetic background. We present data on gene expression and juvenile hormone levels from young and old bees, from both solitary and social nests. We found 940 genes to be differentially expressed with age in solitary females, *versus* only 14 genes in social dominant females, and seven genes in subordinate females. We performed a weighted gene co-expression network analysis to further highlight candidate genes related to ageing in this species. Primary “ageing gene” candidates were related to protein synthesis, gene expression, immunity and venom production. Remarkably, juvenile hormone titres did not vary with age or social status. These results represent an important step in understanding the proximate mechanisms underlying the remodeling of the fecundity/longevity trade-off that accompanies the evolutionary transition from solitary life to eusociality.

**Significance statement:** The remarkably long lifespan of the queens of eusocial insects despite their high reproductive output suggests that they are not subject to the widespread trade-off between fecundity and longevity that governs solitary animal life histories, yet surprisingly little is known of the molecular mechanisms underpinning their longevity. Using a socially plastic bee in which some individuals of a population are social whilst others are solitary, we identified hundreds of candidate genes and related gene networks that are involved in the remoulding of the fecundity/longevity tradeoff. As well as identifying candidate ageing genes, our data suggest that even in incipient stages of sociality there is a marked reprogramming of ageing; long live the queen.

## Introduction

### Ageing and the costs of reproduction

There is longstanding empirical support for the costs of reproduction, and more specifically for the trade-off between fecundity and longevity (Maynard-Smith 1958). Typically, investment in the production and care of offspring has a negative effect on survival (Edward and Chapman 2011). Evidence for such a trade-off has been found in organisms ranging from zebra finches (Alonso-Alvarez et al. 2004) to Columbian ground squirrels (Festa-Bianchet and King 1991), with much support found in model systems such as *Drosophila melanogaster* (Flatt 2011).

Previous studies have helped identify some of the proximate mechanisms underlying this widespread negative correlation between reproduction and lifespan. Endocrine pathways may play a role for instance, with juvenile hormone (JH) correlating positively with reproduction but negatively with lifespan in *Drosophila* (Flatt, Tu, and Tatar 2005), in line with the antagonistic pleiotropy theory of ageing (Medawar 1952; Williams 1957). Additionally, oxidative stress, a molecular marker of senescence according to the free radical theory of ageing (Harman 1956), has been proposed as a mechanism mediating the trade-off between reproduction and lifespan, with reproduction leading to a decrease in antioxidant defences in zebra-finches (Alonso-Alvarez et al. 2004), and increased egg production inducing increased susceptibility to oxidative stress in *Drosophila melanogaster* (Wang, Salmon, and Harshman 2001). Another widespread life history mediator is the nutrient-sensing pathway, more specifically the insulin/insulin-like signalling pathway (IIS) and the target of rapamycin (TOR) pathway (Kapahi et al. 2010). Indeed, these pathways mediate the link between reproductive investment in many organisms, including *D. melanogaster* (Flatt et al. 2008), *Caenorhabditis elegans* (Kenyon 2010), and humans (Blagosklonny 2010).

While evidence for a trade-off between reproduction and longevity is abundant, controversy remains. Germ-line ablation does not cause any difference in lifespan in *D. melanogaster* males, and even slightly decreases lifespan in females (Barnes et al. 2006). The insulin/IGF1 pathway, previously thought central in mediating the fecundity/longevity trade-off across animals, has been shown to control longevity and fecundity independently of each other in *C. elegans* (Dillin, Crawford, and Kenyon 2002).

### Remoulding of the fecundity/longevity trade-off in eusocial insects

Adding to this controversy, a major challenge to the existence of a universal fecundity/longevity trade-off is the remarkable case of eusocial insects, where queens are often the only reproductively active individuals in their colony, yet also live up to 30 times longer than their worker counterparts (Carey 2001). Reproductive individuals in species exhibiting complex eusocial behaviour such as honeybees and ants, but also hemimetabolous insects like termites, thus do not exhibit a negative correlation between reproduction and lifespan (Page and Peng 2001). This remoulding of the expected longevity/fecundity trade-off in eusocial insects has also been supported experimentally. In the ant *Cardiocondyla obscurior*, mating appears to incur no cost in terms of longevity, as mated queens live longer than virgin queens (Schrempf, Heinze, and Cremer 2005), and enforced changes in egg-laying rate do not affect the longevity of queens (Schrempf et al. 2017). Moreover, in honeybees, workers which develop under queenless conditions have higher reproductive potential and also live longer than workers developing in queenright colonies, thus seemingly circumventing the trade-off between fecundity and longevity (Kuszewska et al. 2017).

Several molecular mechanisms have been suggested to underlie the apparent remoulding of the fecundity/longevity trade-off in insects exhibiting complex eusocial behaviour compared to solitary insects and other solitary organisms. For instance, caste-specific differences in somatic maintenance have been observed in ants, with *Lasius niger* queens exhibiting higher expression of somatic repair genes than workers (Lucas, Privman, and Keller 2016). Evidence suggests that a rewiring of central endocrine pathways may underlie the remoulding of the trade-off. Specifically, titres of JH and vitellogenin (a yolk precursor) are both positively correlated with reproduction at the expense of longevity in solitary insects, but the JH/vitellogenin network connectivity varies in social insects (Rodrigues and Flatt 2016). In *Apis mellifera* for instance, JH and vitellogenin are not connected in queens, as JH titres are low throughout their adult life, while vitellogenin levels increase rapidly as they start to lay eggs and remain at high levels as they age (Hartfelder and Engels 1998). In workers, JH and vitellogenin are even negatively connected, and the increase in JH levels once workers become foragers actually sets a limit on their adult lifespan by promoting immunosenescence (Amdam et al. 2005). Hence, in *A. mellifera* queens and workers, vitellogenin is seemingly positively related to somatic maintenance and thus longevity, with high vitellogenin levels making them more resistant to oxidative stress (Seehuus et al. 2006; Corona et al. 2007). By circumventing the positive correlation between JH and vitellogenin, and given the positive impacts of vitellogenin on longevity in this species, honeybee queens are thus able to maintain high reproductive function without sacrificing longevity.

Recently, de-regulation of transposable elements (TEs), which has previously been linked to ageing in many species (De Cecco et al. 2013; Li et al. 2013), has also been suggested as a mechanism allowing a positive correlation between reproduction and lifespan in other eusocial insect lineages: in the termite *Macrotermes bellicosus*, TE-silencing pathways are differentially expressed between sterile and reproductive castes, possibly underlying the striking differences in lifespan (Elsner, Meusemann, and Korb 2018). Although a multitude of mechanisms that promote queen longevity have been proposed in eusocial insects, we still lack a clear understanding of when and how this apparent remoulding of the fecundity/longevity trade-off evolved (Toth, Sumner, and Jeanne 2016). At least within Hymenoptera, eusociality along with a remoulding of a fecundity/longevity trade-off has been suggested to have evolved independently at least eight times (Peters et al. 2017).

### Investigating the remoulding of the fecundity/longevity trade-off in the socially polymorphic orchid bee *Euglossa viridissima*

Studies have investigated the fecundity/longevity trade-off in insect species across the sociality gradient, in particular primitively social and socially polymorphic species. The reversal of the trade-off is evident in primitively social wasps (Toth, Sumner, and Jeanne 2016), shedding light on the mechanisms at play during early stages of social evolution within wasps, such as caste-specific modulation of reproduction by JH (Kapheim 2017). Socially polymorphic species such as the hover wasp *Parischnogaster alternata* (Bolton et al. 2006), the sweat bee *Megalopta genalis* (Kapheim et al. 2013), or the orchid bee *Euglossa viridissima* represent excellent study organisms for such investigations; the behavioural plasticity known for these species allows a direct comparison of solitary and social individuals within the same species, or even within the same population. Such comparisons allow us to identify molecular mechanisms underlying the remoulding of the trade-off in early stages of social evolution, thus distinguishing them from mechanisms which maintain or reinforce this remoulding in complex insect societies (Séguret, Bernadou, and Paxton 2016; Shell and Rehan 2018).

The Neotropical orchid bee *E. viridissima* Friese, 1899, is facultatively eusocial, with solitary and social nests co-occurring within the same population (Cocom-Pech et al. 2008; de May-Itzá et al. 2014). All nests established by a “foundress” female are initially solitary until the first brood emerges approximately two months after founding. Nests are commonly reused for a second brood, and can be reactivated as a single-female nest (solitary) or a multi-female nest in which daughters from the first brood remain in the natal nest and share in care of future broods (social nests). In such multi-female nests, a dominance hierarchy is established through aggressive behaviour, with a reproductive skew in favor of the mother (Cocom-Pech et al. 2008).

In this study we aimed to determine how social organisation and reproductive status (caste) influence lifespan in *E. viridissima* females. Specifically, we investigated changes in gene expression and JH titre with age in reproductive individuals from solitary nests, and compared these to observed changes with age in dominant and subordinate females from social nests. In particular, we aimed to identify candidate genes and gene networks that are linked to differential ageing in relation to reproductive and social status, *i.e*. genes potentially connected with fecundity and longevity. Such gene candidates identified within a socially polymorphic species are central to our understanding of the reversal of the fecundity/longevity trade-off in eusocial insects.

## Materials and Methods

### Experimental setup and sample collection

All *E. viridissima* samples were collected at the Department of Apiculture of the Campus of Biological Sciences and Animal Husbandry, Autonomous University of Yucatán in Xmatkuil, Mexico (89.37°W, 20.52°N, sample collection permit n° 41593). Wooden boxes (7 × 3 × 3 cm) with an inner coating of beeswax and stingless bee cerumen were placed around the campus, with a glass cover between the box and wooden lid to facilitate observations. Females from the wild observed constructing cells in or bringing back provisions to a nest box were individually marked on the thorax with a diamond tipped pen. Nests were checked three times weekly from February 2016 until June 2018 to record the presence of marked females. In multifemale nests, the hierarchy of the females (dominant, subordinate) was determined by observing nests until individuals could be classified based on behaviour (Supplementary Table S1). For multifemale nests with only two females, observations continued until a female was observed returning to the nest with pollen, which characterised this female as the subordinate. In multifemale nests with three or more females, the individual spending most time in the nest, on the brood cells, and exhibiting the largest number of aggressive behaviours towards other females was characterised as dominant (Boff et al. 2015).

For females from solitary nests, and dominant females from social nests, both young (< 1.5 months since marking) and old (> 1.5 months since marking) individuals were collected. The establishment of a threshold of 1.5 months was based on personal observations in the field and on general life history: we estimated lifespan in *E. viridissima* to be of 2-4 months based on anecdotal observations of *E. viridissima* during previous studies of this species at the same campus sites and on what is known for the socially polymorphic *Euglossa melanotricha* (Andrade-Silva and Nascimento 2015). Given that all nests are first established as solitary, only becoming social during the founding female’s second brood phase if her daughters from the first brood remained in the nest (Cocom-Pech et al. 2008), dominant females were, by definition, older than the age threshold set in this study, even during early stages of social life. However, in some cases, dominant females died or left the nest, and one of their subordinate daughters took over as the dominant female. These females were below the set age threshold and were therefore sampled here as young dominant females. Since “worker” (subordinate female) lifespan was unknown for this species at the time of sampling, but is generally much shorter than queen lifespan in other eusocial hymenopteran species (Carey 2001; Séguret, Bernadou, and Paxton 2016), we used three weeks as a threshold for distinguishing “young” *vs* “old” workers based on previous personal observations of this species at the same sites, and supported post-hoc by comparison of transcriptomes (see Results, Figure 1). Collection continued until at least five individuals were collected for each combination of age and social status, except for old subordinate females which were rarely found in the field.

**Fig. 1.**
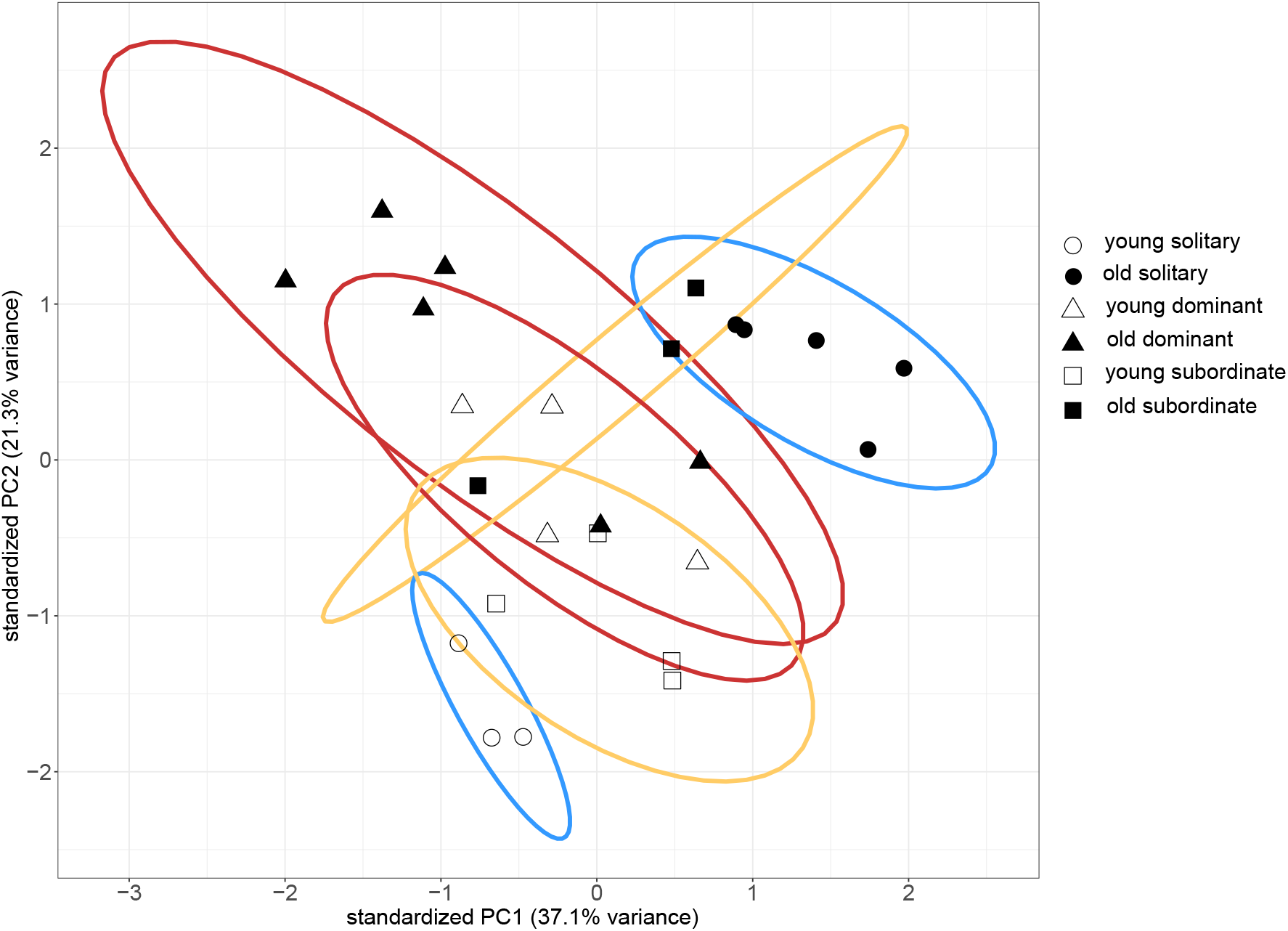
Principal component analysis (PCA) of variance-stabilised RNA read counts of young and old females from solitary and social nests. Each point represents the expression profile of one individual across the 958 genes which were differentially expressed between young and old individuals, cumulative across all social types (solitary: 940 DEGs, dominant: 14 DEGs, subordinate: 7 DEGs, with 2 DEGs shared between solitary and subordinate, and 1 DEG shared between solitary and dominant, see Fig. 2). Axis labels indicate the amount of variance in gene expression explained by the first two principal components (PC1 and PC2). Ellipses represent 95% confidence levels, and further illustrate the social type (blue=solitary; red=dominant; yellow=subordinate) of individuals.

Upon collection, each individual was weighed and its thorax width (intertegular distance) was measured using a Canon EOS 60D camera and the imaging software Motic Images Multifocus Pro 1.0. Next, a haemolymph sample (1.5 - 4 μL) was collected from the abdomen using microcapillaries for JH measurements. Each haemolymph sample was transferred to Teflon capped GC-vials containing 500 μL acetonitrile that was then stored at −80°C. Whole bees were then kept overnight at −80°C, and once frozen, the abdomen was removed and transferred to RNAlater^®^ (Sigma-Aldrich) for storage at −80°C. Altogether, 34 individuals were used for molecular analyses (Supplementary Table S2).

### RNA extraction and sequencing

Total RNA was extracted from each individual (whole abdomen) using the RNeasy^®^ Plus Mini kit, including DNase I digestion, according to manufacturer’s instructions, using 350 μl starting material (Qiagen, Hilden, Germany). RNA concentration, purity and integrity were measured using an Agilent 4200 TapeStation and 6 RNA ScreenTapes. Thirty-two samples which passed our quality criteria (260/280 = 2.1 ± 0.1, RINe > 9, total RNA mass > 1 μg) were used for RNA sequencing (Supplementary Table S2). Library preparation and transcriptome sequencing were undertaken at Beijing Genomics Institute (BGI, Shenzhen). For each sample, a cDNA library was prepared with the TruSeq RNA Library Preparation kit (Illumina) and SuperScript II Reverse Transcriptase (Thermo Fisher Scientific). Libraries were sequenced on an Illumina HiSeq4000 platform (11 samples per lane) to generate 100 bp paired-end reads, with ~4 GB of raw data per sample, *i.e*. 29 million read pairs per sample on average (range 24-35 million).

Raw reads were quality-checked with FastQC v0.11.5 (Andrews 2010), then filtered and trimmed using Skewer v0.2.2 (Jiang et al. 2014) to remove low-quality bases and reads, adapter contamination and reads shorter than 70 bp.

#### Exclusion of individuals from a sister species

Individuals belonging to the cryptic sister species *Euglossa dilemma* were identified through sequence comparison of the olfactory receptor gene *or41*, which has been described as a means of differentiating between *E. viridissima* and *E. dilemma* (Brand et al. 2020) (Supplementary Table S3). Six *E. dilemma* individuals were identified and subsequently excluded from further analyses in order to avoid any bias, although overall patterns in the data did not change when including these individuals (Supplementary Figure S4 and Tables S6, S7).

### Differential gene expression analysis

#### Transcriptome assembly, genome annotation, read mapping and quantification

For full details on the transcriptome assembly and genome annotation, as well as software references, see Supplementary File S8. The transcriptomic analyses are based on the previously published draft genome sequence assembly for *E. dilemma* GCA_002201625.1 (Brand et al. 2017). Repeats were soft-masked (35.30% of the total genome assembly length) using bedtools v2.27.1 based on repeat annotations from Tandem Repeats Finder v4.09 and RepeatMasker. To improve the previously published annotation of this genome, which was based solely on gene predictions and homology to *Apis mellifera* proteins (Brand et al. 2017), we used Funannotate v1.5.1 with the previous gene annotation (edil. 1.0.annotations.gff), novel experimental evidence derived from our RNAseq data generated in this study, namely: (i) RNAseq paired-end reads aligned to the sister species *E. dilemma* genome with HiSat2 v2.1.0, (ii) a transcriptome assembly with binpacker v1.1 from one sample, and (iii) a genome-guided transcriptome assembly from four samples with Stringtie2 v1.3.3b, as well as data from published studies such as (iv) an RNAseq data-based transcriptome assembly of *E. dilemma* using Trinity v1.5.1, and (v) transcripts of the closely related orchid bee *Eufriesea mexicana* (GCF_001483705.1). Further, protein sequences from five related bee species (*Bombus impatiens:* GCF_000188095.2; *B. terrestris*: GCF_000214255.1; *Apis mellifera:* GCF_003254395.2; *Melipona quadrifasciata:* GCA_001276565.1; *Eufriesea mexicana:* GCF_001483705.1) and Uniprot (sprot) were used for homology-based evidence. Additionally, genes were predicted from the *E. dilemma* genome using SNAP v2006-07-28 based on SNAP’s *Apis mellifera* HMM dataset, Genemark-ET, Augustus v3.3 based on BUSCO3 v3.0.2 training, and the PASA pipeline v2.3.3.

In brief, PASA was used to train gene predictions which were then performed using Genemark-ES, PASA, Augustus v.3.3, and CodingQuarry v2.0. Predictions as well as additional, previous annotations were then integrated in Evidence Modeler v.1.1.1. Too short, gap-spanning or repeat-overlapping gene models were removed and tRNA genes were detected with tRNAscan SE v2.0.0. Previous gene models were then updated with PASA and Transdecoder v5.5.0 based on RNAseq evidence and expression levels of models at each locus. Genes were functionally annotated using PFAM v32.0, the UniProt database v2018_11, EggNog Annotations (eggnog_4.5/hmmdb databases: Arthropoda, Insecta, Hymenoptera, Drosophila), MEROPS v12.0, CAZYmes in dbCAN v7.0, BUSCO Hymenoptera models v3.0.2, Hymenoptera odb9, SignalP v4.1, and InterProScan5 v5.33-72.0. After processing and quality filtering, gene models were compared to the previous *E. dilemma* annotation (v1.0). Non-duplicated gene models from Edil1.0 which were missing in the Funannotate annotations (largely due to missing stop codons (n=268), gap spanning (n=16), or other (n=13)) were appended. The final annotation contained 24,991 gene models (including 126 tRNA genes), and was estimated to be 80.6% complete (BUSCO3: 3,560 were found out of 4,415 single copy protein coding orthologs conserved across Hymenoptera, and 89 (2.0%) of these were duplicated).

The set of 24,991 gene models was then used for subsequent transcriptome analyses. Trimmed and filtered RNAseq reads were mapped to these gene models and were simultaneously quantified using Kallisto v0.46.2 (Bray et al. 2016).

#### Differential expression analyses

In order to analyse differential gene expression between young and old individuals within each group (see below), we used the tximport package v1.12.3 (Soneson, Love, and Robinson 2015) in R v3.6.1 (R Core Team 2019) to import transcript-level read quantification data, and convert these to gene-level quantification values (Supplementary Table S9). We then ran DESeq2 v1.18.1 (Love, Huber, and Anders 2014) with default settings (adjusted *p*-value calculated to correct for multiple testing following the BH method (Benjamini and Hochberg 1995)) to perform pairwise comparisons of gene expression profiles between young and old individuals of a given social status (*i. e*., solitary, social dominant or social subordinate), and between individuals of different social status for a given age. Differentially expressed genes (DEGs) were identified as those with a *p*-adjusted value below 0.05. After plotting gene expression profiles of all individuals, excluding *E. dilemma* individuals, in a Principal Component Analysis plot (PCA of variance stabilised read counts, with genes as columns and samples as rows in the PCA matrix, Supplementary Figure S5), individual eug6 was identified as an outlier and was excluded from gene expression analyses. All PCA figures were produced using the R package ggbiplot v0.55 (Vu 2011), with ellipse probabilities (if ellipses shown) set to 95% confidence level (ellipse.prob=0.95). The final dataset comprised 25 *E. viridissima* transcriptomes.

To identify the genes for which expression levels were affected by the interaction between age and social status, we performed a likelihood ratio test to contrast the following models in DESeq2: Full model (age + social_status + age:social_status), Reduced model (age + social_status). This contrast identified genes with a significant change in expression associated with the interaction of the two factors, age and social status.

### Weighted gene co-expression network analysis

#### Network construction and correlation of modules with phenotypic traits of interest

Weighted gene co-expression network analysis was carried out using the R package WGCNA (Langfelder and Horvath 2008). The aims of this analysis were to (a) identify sets of co-expressed genes (modules), (b) calculate module eigengenes (*i.e*. values representative of the gene expression profile in a module) and correlate these with phenotypes of interest (here, social status and age), and finally (c) identify “hub genes”, *i.e*. genes that were most highly connected, within modules which were significantly associated with the phenotypic trait of interest (here, age). Additionally, the network analysis was performed to identify age-related hub genes overlapping with those differentially expressed between young and old individuals. For this analysis, genes with read counts lower than 10 in 90% of the samples were removed, and raw read counts were transformed using the varianceStabilizingTransformation function in DESeq2, as recommended by Langfelder and Horvath (2008). The input table thus consisted of a matrix of 25 individuals across all social types and ages and 12,902 gene expression values (Supplementary Table S10). Modules of co-expressed genes were defined using average linkage hierarchical clustering with the topological overlap-based dissimilarity measure, with parameters set according to recommendations of Zhang and Horvath (2005) and Langfelder and Horvath (2008) (soft thresholding power set to 14 as the lowest value for which scale-free topology fit index reached 0.9; minimum module size set to 30 genes; modules of highly co-expressed genes merged using a cut-off value of 0.2).

#### Hub genes correlated with age and social status

Module eigengenes were correlated with two phenotypic traits: social status and age. Hub genes from each module associated with age, *i. e*. genes which were most highly connected within those modules (intramodular connectivity > 0.75) were extracted and concatenated to create a list of hub genes associated with age.

### Overlap of age-related genes from two independent analyses

Genes found to be common to the list of hub genes associated with age in the WGCNA and the list of genes found to be differentially expressed with age were considered to be of particular interest. The functions of these overlapping genes were further investigated through orthology analyses (see below).

### Functional annotation

#### Selection of “top genes” from gene lists of interest

Functional annotation was done for top genes from each gene list of interest. For lists of significant DEGs, the top ten genes, *i. e*. genes with the highest absolute log2 fold change, were extracted from the lists of 1) genes up-regulated in young compared to old, and 2) genes up-regulated in old compared to young, separately for a) solitary, b) dominant and c) subordinate females (six lists in total). The top ten genes whose expression was affected by the interaction between age and social status were defined as genes with the highest absolute log2 fold change value in the model contrast (likelihood ratio test). For hub genes from the modules associated with age in the network analysis, the top ten genes were defined for each module as those with the highest intramodular connectivity.

Finally, all genes found to be associated with age in both the differential expression analysis and the network analysis were considered of particular interest and were functionally annotated.

#### *Assigning gene functions based on orthology with* A. mellifera and Bombus *spp*

To assign functions to genes highlighted as relevant with regards to ageing, we searched for the known function of their closest ortholog. OrthoFinder v2.3.7 (Emms and Kelly 2015) was run with default settings on protein level using the genome annotation we produced for *E. dilemma* and protein sets of three closely related species with well-annotated genomes: *A. mellifera* (GCF_003254395.2), *Bombus terrestris* (GCF_000214255.1), and *Bombus impatiens* (GCF_000188095.3). For all top gene lists, ortholog protein names were extracted from *A. mellifera* annotations if available as this is the most extensively annotated genome, or from the annotations in *Bombus* spp. if no *Apis* ortholog was found.

For the remaining 19% of *Euglossa* top genes which did not have an ortholog in any of these three species (see Results), nucleotide sequences were searched against the NCBI nucleotide database (NCBI, database downloaded on 07.01.2020) using BLAST v2.7.1 (Altschul et al. 1990). The closest orthologous gene was then determined based on its best hit according to its lowest e-value.

Ortholog protein names were then searched manually in UniProt for existing functional annotations in the species in which the ortholog was found, or in a closely related species if available. Based on the functional annotations found, each gene was then assigned to one of the following functional categories: energy metabolism, stress response, growth/trophic factors, cell cycle, signalling, venom, protein turnover, gene regulation or TE suppression.

### Deposition of genome annotation files and raw sequence reads

All raw sequence reads for each sample, including *E. dilemma* individuals, have been deposited in the NCBI Sequence Read Archive under Bioproject ID PRJNA636137. The updated genome annotation for *E. dilemma* has been deposited in Dryad (see Data Availability Statement).

### Juvenile hormone quantification and analysis

The same individuals used for the transcriptomic analyses were also used to assess juvenile hormone (JH) levels in the haemolymph by a radioimmunoassay. The protocols for JH extraction from the haemolymph samples in acetonitrile and preparation of the samples in the radioimmunoassay (RIA) followed the detailed description for JH quantification in the honeybee (Hartfelder et al. 2013) and wasps (Kelstrup et al. 2018), using a JH-specific antiserum, tritiated [10-^3^H(N)]-JH III (specific activity 19.4 Ci/nmol, Perkin Elmer Life Sciences, Waltham, MA, USA), and synthetic juvenile hormone-III (Fluka, Munich, Germany) as the non-radioactive competitor. For the haemolymph JH titre calculations, we used a non-linear four-parameter regression. After exclusion of *E. dilemma* individuals, 21 *E. viridissima* individuals were included in the analysis (Supplementary File S2).

All statistical analyses were performed in R (v3.6.1). To investigate the effect of age and social status on JH titres, we performed a generalised linear mixed model on our log-transformed data using the package lme4 v1.1-21 (Bates et al. 2015). A stepwise Akaike Information Criterion (AIC) method for variable selection was applied, with the best-fitting model including age, social status and the interaction between age and social status as fixed factors, and nest ID and date of collection as random factors. An ANOVA was then used to test for significant effects.

## Results

### RNA sequencing

After trimming and filtering, sequenced libraries contained on average 29 million read pairs (range ± SD = 24-35 million reads ± 5 million read pairs), of which 79% (range 66-79 ± 19% SD) could be mapped to the *E. dilemma* genome annotation.

### Gene expression changes with age in solitary but not in social females

A total of 19,136 genes had at least one read count, representing 77% of annotated genes (for a PCA across all these genes, not just those which were differentially expressed, see supplementary Figure S12). The PCA of variance-stabilised read counts across genes which were differentially expressed between young and old individuals, cumulative across all social types (Figure 1) showed a clear distinction between the gene expression profiles of young and old solitary females across both PC1 (explaining 37% of the variance) and PC2 (explaining 21% of the variance). Young and old subordinate females from social nests also exhibited distinct gene expression profiles, although to a lesser extent than solitary females. In dominant females, however, gene expression profiles did not show a clear pattern with respect to age; data points from young and old individuals overlapped (Figure 1).

Differential gene expression analyses revealed 940 genes to be significantly differentially expressed with age in solitary females, versus only 14 genes differentially expressed with age in dominant females, and seven in subordinate females (adjusted *p* < 0.05, Supplementary File S14 and Figure 2a). Additionally, solitary and dominant females exhibited increasingly divergent expression profiles with age, with 213 DEGs between young solitary and young dominant females, versus 2,601 DEGs between old solitary and old dominant females. Subordinate females from social nests showed gene expression profiles which were intermediate between those of solitary and dominant females, with 38 DEGs between young solitary and young subordinate females, 53 DEGs between old solitary and old subordinate females, 212 DEGs between young subordinate and young dominant females, and 38 DEGs between old subordinate and old dominant females (for a list of differentially expressed genes in each comparison, see Supplementary File S14). Finally, old dominant females exhibited considerably more differences to young subordinate females compared to young dominant females (1,007 and 14 DEGs, respectively), despite young dominant females in our study having been born as subordinate females which switched to the dominant position upon their mother’s absence from the nest (see Methods).

**Fig. 2.**
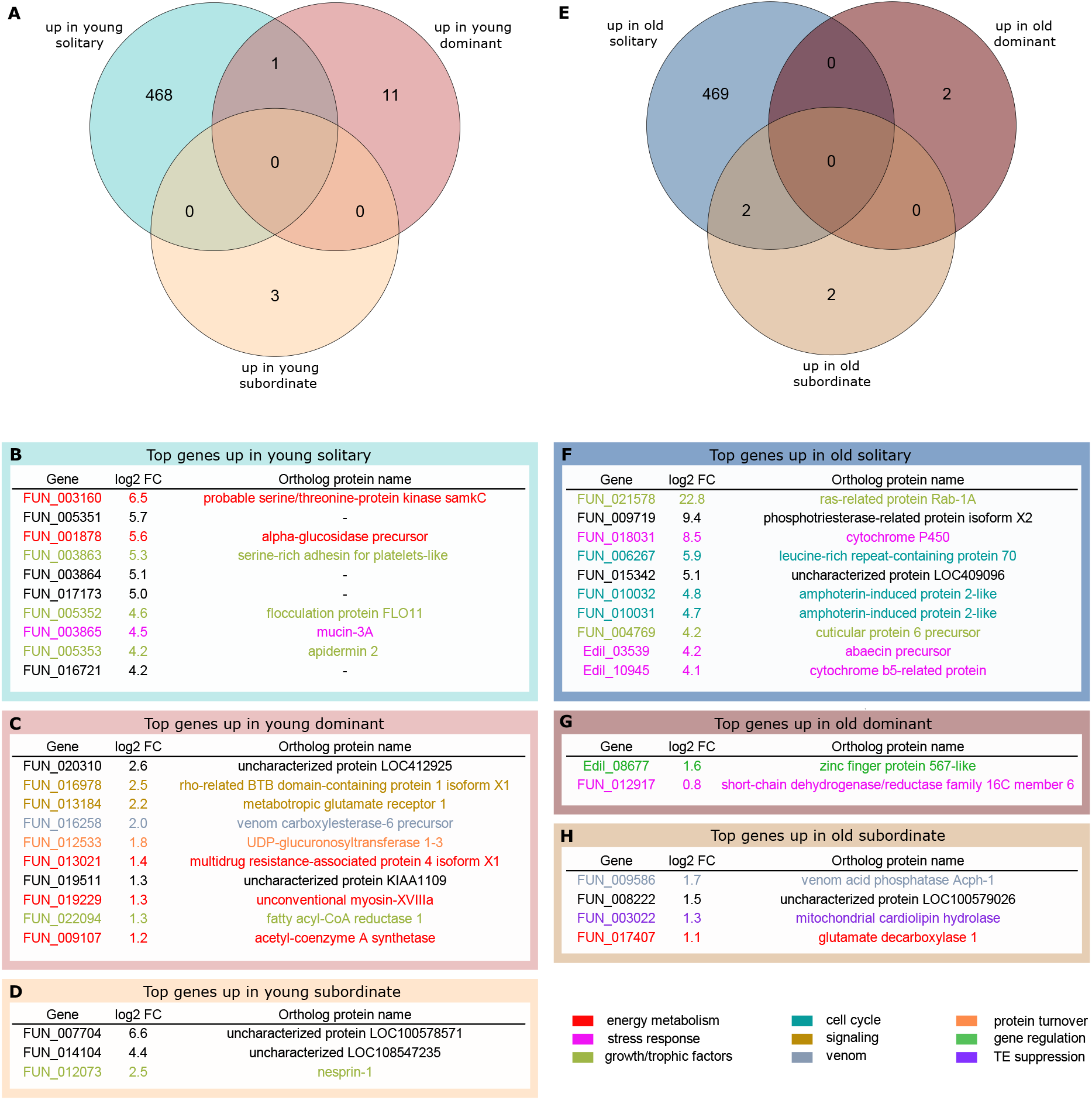
Venn diagrams and ortholog protein names of genes differentially expressed with age in solitary, dominant and subordinate females. The Venn diagram and tables on the left (A-D) represent genes upregulated in young compared to old individuals. The Venn diagram and tables on the right (E-H) represent genes upregulated in old compared to young individuals. Venn diagrams and table frames are colour-coded according to comparison as in Figure 1. Tables B-D and F-H show, for each comparison, the list of top genes (highest absolute log2 fold-change) as well as the protein names of their closest found orthologs. The listed genes are colour-coded according to putative functional category based on annotations in UniProt (see legend, bottom-right). Genes left in black were not assigned.

Functional annotation of the top ten genes upregulated in young compared to old solitary females revealed several genes related to metabolic pathways (*e.g*. alpha-glucosidase precursor, generally involved in the break-down of starch molecules into glucose), as well as genes involved in stress response and in growth and trophic factors (Figure 2b; for a full table of functional annotations see Supplementary File S15). Top genes upregulated in old compared to young solitary females were mainly involved in the stress response (*e.g*. a gene from the cytochrome P450 superfamily, which plays a role in protection against reactive oxygen species in *Apis cerana cerana* (Zhu et al. 2016)) and in the cell cycle, as well as in growth and trophic factors (Figure 2f). Top genes upregulated in young *vs*. old dominant females were involved in a range of pathways, from signalling (*e.g*. metabotropic glutamate receptor 1, a G-protein coupled receptor thought to play a role in the action of glutamate in the central nervous system of *Mus musculus* (Charette et al. 2016)) to energy metabolism (*e.g*. unconventional myosin-XVIIIa, putatively involved in motor activity and ATP binding (Redowicz 2007)) (Figure 2c). Genes upregulated in old compared to young dominant females were involved in gene regulation (*e.g*. zinc finger protein 567-like, likely involved in the regulation of transcription (Fasken, Corbett, and Stewart 2019)) and response to stress (Figure 2g). Finally, genes upregulated in young compared to old subordinate females were mainly uncharacterised, but one was linked to growth and trophic factors (nesprin-1, potentially involved in subcellular spatial organisation (Zhang et al. 2001)) (Figure 2e). Genes upregulated in old compared to young subordinate females related to the silencing of transposable elements (mitochondrial cardiolipin hydrolase (Todeschini et al. 2010)), energy metabolism and venom production (Figure 2h). The one gene which was upregulated in young compared to old individuals in both solitary and dominant females is to date uncharacterised (Gene ID FUN_020310). The two genes that were convergently upregulated with age in solitary and subordinate females were related to TE suppression and energy metabolism (Gene IDs FUN_003022 and FUN_017407, Supplementary File S15).

By contrasting two models in DESeq2 (with *versus* without the interaction between age and social status), we detected 314 genes exhibiting different changes in expression with age according to social status (likelihood-ratio test adjusted *p* < 0.05, Supplementary File S14). Functional annotation of these genes revealed an overlap with functions of genes differentially expressed between young and old solitary and subordinate females, including links to the stress response, metabolic pathways, growth and trophic factors, and the cell cycle (Supplementary File S15).

### Modules of co-expressed genes correlated with age

Independent identification of genes for which expression patterns correlate with age was performed using the same gene expression dataset, following the Weighted Gene Correlation Network Analysis approach (WGCNA (Langfelder and Horvath 2008)). A total of 10,446 genes (81% of the 12,902 expressed genes used in this analysis, see Methods and Supplementary File S10) were assigned to 26 modules forming a co-expression network. Correlation of each module’s eigengene with age and social status revealed that four modules of co-expressed genes were significantly associated with age, 10 modules were associated with the solitary phenotype regardless of age, 13 modules were associated with the dominant phenotype (seven of which overlapped with those associated with solitary phenotype, Supplementary File S13 and Supplementary Figure S16), and three modules were associated with the subordinate phenotype, again irrespective of age. However, these three latter modules were also associated with the dominant phenotype (Supplementary Figure S16).

For the four modules significantly associated with age, we extracted hub genes, *i.e*. genes with an intramodular connectivity > 0.75 (intramodular connectivity range 0-1). This resulted in a total of 35 genes (out of 2,053 genes in total belonging to the four age-correlated modules) revealed to be significantly related to age (Figure 3a; for the full list of genes see Supplementary File S13, and for functional annotation of the top ten hub genes in each module see Supplementary File S15). Functions of hub genes from the four age-related modules mainly related to protein synthesis, energy metabolism, fatty acid biosynthesis, and response to stress (Figure 3).

**Fig. 3.**
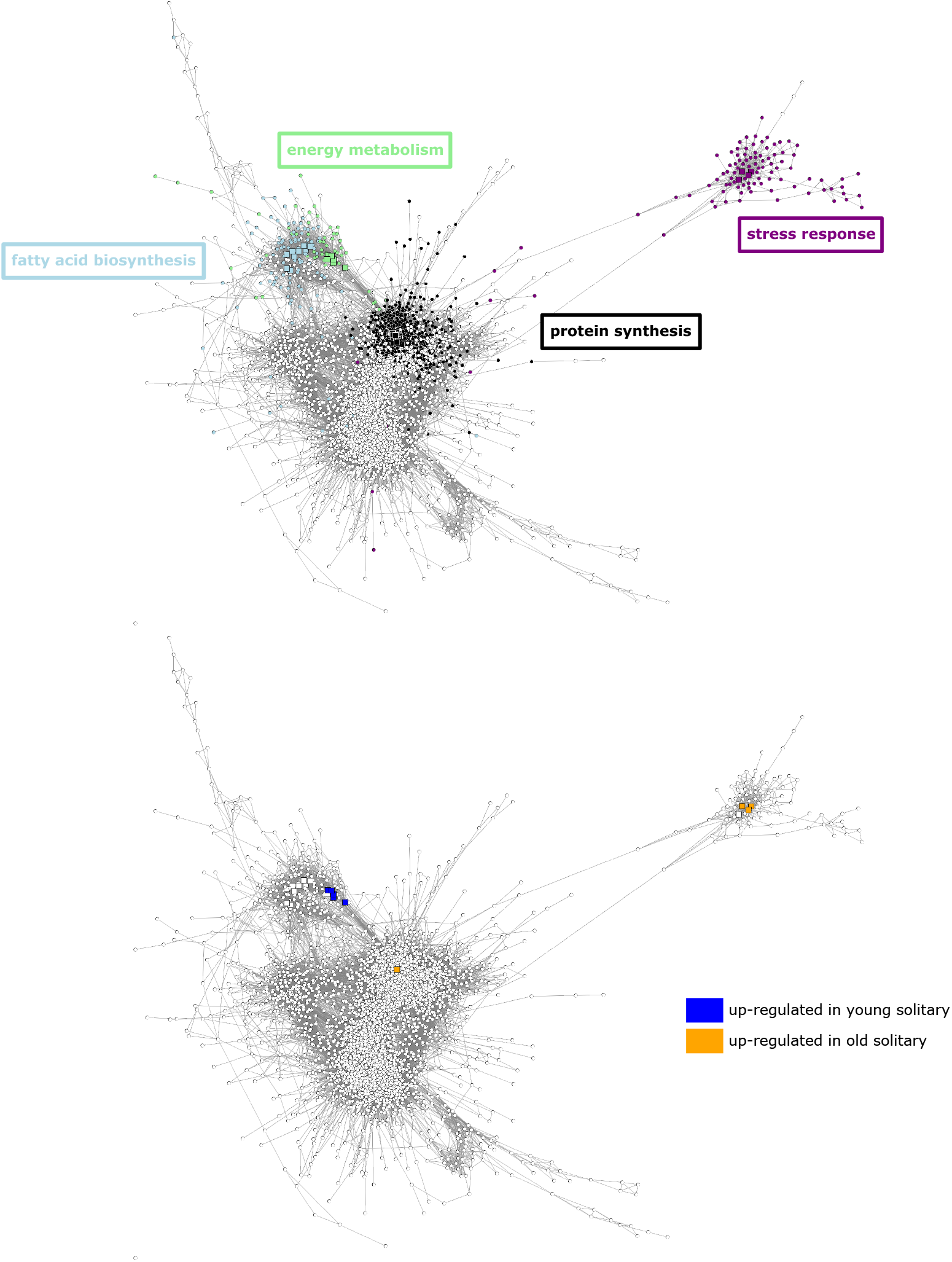
Weighted gene co-expression networks across all expressed genes. Coloured nodes in A represent all genes from the four age-associated modules (purple, black, blue, green), with each colour representing a different module. Square nodes represent hub genes (intramodular connectivity > 0.75). Framed words indicate the top functions of the top ten hub (most highly connected) genes in each module. In B, coloured nodes represent genes from modules in A which are differentially expressed between young and old solitary females (blue = up-regulated in young, orange = upregulated in old). Coloured nodes in network B thus highlight the overlap between the genes found to be significantly associated with age in each independent analysis, i.e. the coexpression network analysis and differential gene expression analysis. A reduced version of the network is presented here for visualisation purposes, with 6,105 nodes and circa 500,000 edges.

### Overlap of age-related genes between the differential gene expression and the network analyses

Of the 35 genes for which expression was significantly correlated with age in the WGCNA, ten were also found to be significantly differentially expressed between young and old solitary females (Figure 3b, Figure 4). There was no overlap of age-related genes between the two analyses for dominant or subordinate females. The overlap for solitary females was significantly higher than expected by chance (hypergeometric test, *p* < 0.001). Functional annotation of these overlapping genes revealed that these included genes related to immunity and stress response (*e.g*. serine protease inhibitor genes 27A and 88Ea, which negatively regulate the melanisation cascade and the Toll signalling pathway in *Drosophila melanogaster* (Tang et al. 2006)), as well as to gene translation and protein turnover (*e.g*. 60S ribosomal protein L34 is potentially involved in translation in *Apis cerana*, and kynurenine/alpha-aminoadipate aminotransferase is involved in the synthesis of amino acids), and to venom production (venom acid phosphatase Acph-1) (Figure 4b).

**Fig. 4.**
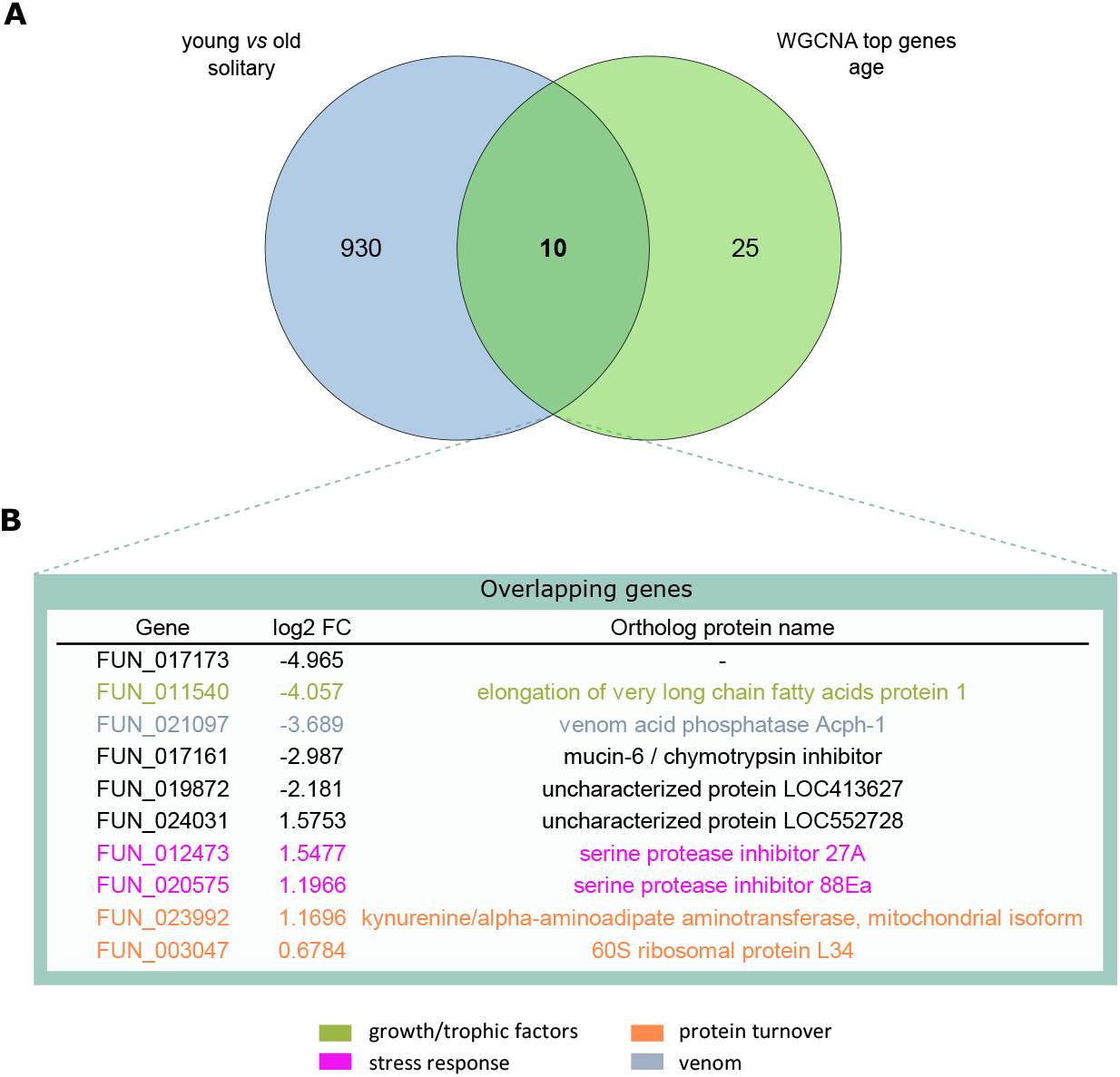
Venn diagram (A) and functional annotation (B) for genes significantly associated with age in solitary females in the WGCNA network and that are differentially expressed between young and old solitary females. The green circle in A represents the cumulative 35 hub genes (intramodular connectivity > 0.75) across all four age-related modules in the WGCNA. The blue circle shows the number of DEGs between young and old solitary females. Table B shows the functional annotation of the ten genes shown in bold (intersection in A), which were found to be significantly associated with age in solitary females in both the DEG and the WGCN analyses. Genes with a negative log2 fold change were up-regulated in young compared to old solitary females; genes with a positive log2 fold change were up-regulated in old compared to young solitary females. The listed genes are color-coded according to functional category.

### No change in juvenile hormone titres with age or social status

After controlling for nest of origin and sampling date, there were no significant differences in juvenile hormone titre in relation to age, social status, or the interaction between the two factors (ANOVA, age: χ2(1,21) = 0.01, *p* = 0.92; social status: χ^2^(2,21)=4.75, *p* = 0.09; age × social status: χ2(1,21) = 0.68, *p* = 0.41; Supplementary Figures S17, S18).

## Discussion

### Differential gene expression with age depends on social status

#### Solitary females exhibit extensive changes in gene expression profiles with age

The difference in gene expression profiles of young and old solitary females, and the high number of differentially expressed genes between these two groups (940 genes) strongly suggest that solitary females undergo substantial physiological changes with age. Functional annotation of these 940 DEGs revealed that metabolic pathways likely play a major role in (or are affected by) these changes. Genes relating to metabolic activities appear to be upregulated in young rather than old solitary females, suggesting that high metabolic activity early in life may increase senescence, as predicted by the free radical theory of ageing (Harman 1982). In the cryptic sister species *Euglossa dilemma*, young foundress females indeed spend more time foraging for brood provisions, a metabolically costly task, compared to when they later enter the “guard phase”, during which they stay in the nest, guarding their brood cells (Saleh and Ramírez 2019). Interestingly, genes directly related to the oxidative metabolism of various substrates, notably lipids (cytochrome P450, cytochrome b5) are upregulated with age in solitary females. In mammals, cytochrome P450 genes are controlled through tight regulation of gene transcription, and a dysregulation of transcription with age can therefore lead to an accumulation of reactive oxygen species and increased oxidative stress (Zangar, Davydov, and Verma 2004).

The relationship between high metabolic activity and ageing has long been suspected, as the accumulation of reactive oxygen species, an inevitable by-product of metabolic activity, is thought to accelerate senescence, therefore making it a potential hallmark of biological ageing (Alonso-Alvarez et al. 2004; Monaghan 2014). In solitary insects such as *Drosophila*, gene expression changes during ageing overlap with those induced by increased exposure to oxidative stress, including an increase in cytochrome P450 genes (Landis et al. 2004). The *Drosophila* mutant line *methusaleh* and mutants of the *age-1* gene in *C. elegans* are both longer-lived and more resistant to oxidative stress (Lin, Seroude, and Benzer 1998; Larsen 1993), reinforcing the idea of oxidative stress as a cause of senescence.

However, multiple studies have called this idea into question (Pérez et al. 2009; López-Otín et al. 2013), suggesting that the link between metabolism and biological ageing is not so straightforward. Still, the upregulation of metabolic pathways early in life and of genes related to the oxidative stress response later in life are likely signs of senescence in solitary females.

It should be noted that half of the top ten genes upregulated in young compared to old solitary females do not yet have functional annotations; the functional interpretation of genes differentially expressed with age in solitary females should therefore be interpreted with caution.

#### Social females undergo very little transcriptomic change with age

Although very few genes were differentially expressed with age in social compared to solitary females, those may provide insight into ageing patterns associated with the social phenotype in *E. viridissima*.

##### 1) Metabolic pathways: down in old dominant, up in old subordinate

While genes linked to energy metabolism were upregulated in young compared to old dominant females, the opposite trend was observed in subordinate females (although it is worth noting that two of the three genes upregulated in young subordinate females are to date still uncharacterised). This discrepancy may be explained by the different life histories exhibited by dominant and subordinate females in this species. Indeed, while subordinate females may perform more costly tasks later in their life as they must provision for an increasingly large brood, dominant females may be subject to more metabolically costly tasks at a younger age, as they produce the brood and establish their dominance through aggressive interactions with other females in the nest early in the nesting cycle. Although *E. viridissima* has been reported to be remarkably tolerant towards daughters in the nest (Cocom-Pech et al. 2008), we observed multiple cases of agonistic behaviours from the dominant towards subordinate females, as described in several primitively social species as a way of establishing dominance (Freiria, Garófalo, and Del Lama 2017; Boff, Saito, and Alves-dos-Santos 2017).

Interestingly, a similar gene expression pattern has been found in honeybees, in which genes linked to oxido-reductase activity were upregulated with age in workers (Aumer et al. 2018); in the ant *Temnothorax rugatulus*, where the oxidation-reduction process was the only significantly enriched GO term for genes downregulated with age in queen brains (Negroni, Foitzik, and Feldmeyer 2019); and in the termite *Cryptotermes secundus*, where multiple transcripts related to oxidative stress were upregulated in young compared to old queens and kings. Workers showed the opposite trend, with such transcripts being upregulated in older individuals (Kuhn, Meusemann, and Korb 2019).

##### 2) Silencing of transposable elements

The gene coding for mitochondrial cardiolipin hydrolase, which was one of the two genes upregulated in old subordinate females in our study, is required for the activity of the piRNA pathway in *Drosophila melanogaster* (Todeschini et al. 2010). This pathway is known for its role in silencing transposable elements (TEs). Recently, TEs have been highlighted as a potentially major factor underpinning differential ageing between the queen/king and worker caste in the “higher” termite *Macrotermes bellicosus* (Elsner, Meusemann, and Korb 2018), and in the “lower” termite *Cryptotermes secundus* (Kuhn, Meusemann, and Korb 2019). Given the high repetitive proportion of the *E. viridissima* genome (Brand et al. 2017; annotation Edilemma_v2.2 presented here), TE silencing may play an important role in *Euglossa* bees too, in contrast to highly eusocial bees which have very few active TEs (Kapheim et al. 2015).

##### 3) Gene regulation: an underestimated hallmark of ageing?

Functional annotation of genes differentially expressed between young and old dominant females also suggests that the regulation of gene expression may play a role in ageing. A meta-analysis of age-related gene expression profiles using 27 datasets from mice, rats and humans revealed that the “negative regulation of transcription” was indeed one of the GO categories overrepresented in age-related transcriptional profiles, making it a common signature of ageing across those organisms (de Magalhães, Curado, and Church 2009). Frenk and Houseley (2018) also propose dysregulation of gene expression and mRNA processing with age as a candidate “gene expression hallmark” of cellular ageing. Empirical studies have supported this theory: for example, gene expression correlation decreases with age in mice (Southworth, Owen, and Kim 2009), which can be interpreted as transcriptional noise increasing with age (Bahar et al. 2006). Experimental exposure to oxidative stress increases cell-to-cell variation in gene expression, suggesting that dysregulation of gene expression is a possible mechanism linking DNA damage with ageing, cellular degeneration and death. In our study, upregulation of a gene linked to the regulation of transcription (zinc finger protein 567 (Fasken, Corbett, and Stewart 2019)) with age in dominant females, but not in solitary females, suggests that dominant females may actively counter the dysregulation of gene expression with age, thus supporting differential gene regulation as a potential mechanism for the reduced signs of ageing in social dominant females.

### Does sociality prevent ageing?

The considerable age-related differences in gene expression in solitary females but not in the social females observed in this study (neither dominant nor subordinate) suggest that solitary females undergo more extensive physiological changes with age, particularly when compared to dominant females in social nests. The divergence in observed age-related gene expression changes between solitary and social dominant females despite them belonging to comparable age groups are worth noting and support the idea of the co-evolution of extended longevity and eusociality in its early evolutionary stages (Carey 2001). Further studies on other molecular markers related to ageing (López-Otín et al. 2013) in species along the sociality gradient may help deepen our understanding of this issue.

As subordinate females perform costly tasks such as brood provisioning, and given that workers of obligate eusocial species exhibit considerably shorter lifespan than their queen counterparts (Carey 2001), the absence of major changes in gene expression with age in subordinate *E. viridissima* is somewhat surprising. However, the two genes which were convergently upregulated with age in both solitary and subordinate females provide some insight into the costs of the tasks performed by both solitary and subordinate females in *E. viridissima*. These genes were related to TE suppression and energy metabolism, and are thus potentially related to known ageing pathways (Elsner, Meusemann and Korb 2018). Experimental studies in which putatively costly behaviours, such as brood provisioning, are manipulated may further our understanding of the actual costs of such tasks.

An alternative explanation for the differences in gene expression with age observed in the solitary, but not observed in the social females is a potential shift in behaviour with age in the solitary females. It has been proposed for the sister species *E. dilemma* that transcriptomic shifts throughout the solitary phases of the life cycle may be attributed to a shift in behaviour, as the female goes from an actively foraging foundress to a relatively inactive guard, caring for her brood (Saleh and Ramírez 2019). Similar to the differences observed between young and old solitary females in our study species *E. viridissima*, Saleh and Ramirez (2019) found that metabolic pathways were upregulated in foundresses compared to females in the guarding phase. Additional information on the life cycle and associated shifts in behaviour in *E. viridissima* would therefore allow us to better account for this potential factor, and determine whether the differences observed in our study are due to ageing *per se* as opposed to behavioural repertoire. Another factor which may explain the lack of ageing signs in dominant females is the ingestion of subordinate-laid eggs by dominant females. Video recordings of *E. dilemma* nests revealed that oophagy, whereby the dominant female ingests eggs laid by subordinate females in her nest to replace them with her own, occurs commonly in this species (Saleh and Ramírez 2019). If this behaviour also occurs in the closely related *E. viridissima*, this could constitute a notable nutritional source for dominant females, thereby reducing any trade-off generated by the allocation of limited resources.

### Gene co-expression networks as a tool to further highlight “ageing gene” candidates

#### Using network hub genes to independently identify ageing pathways in E. viridissima

Our gene co-expression network analysis was performed in part to identify age-related genes overlapping with those differentially expressed with age, thus narrowing down the list of genes that potentially play a role in the ageing process. The results from the co-expression network analysis highlighted modules of genes for which expression was correlated with age, independent of social status. Functional annotation of these genes indeed revealed that they were related to well-established age-related pathways such as metabolic processes and the stress response (Landis et al. 2004). Protein synthesis was the main function associated with one of these age-related modules (black module in Figure 4). The expression of ribosomal proteins increases with ageing; however, as proteasome levels decrease with age, they are accompanied by a loss of protein homeostasis, a common feature of ageing (García-Velázquez and Arias 2020). Finally, the top ten hub genes of one of the age-related modules (blue module in Figure 4) were related, among other things, to fatty acid biosynthesis (very-long-chain 3-oxoacyl-CoA reductase and fatty acyl-CoA reductase), a pathway which has been linked to ageing in several species (Reis et al. 2011; Proshkina et al. 2015). Thus, hub genes from age-related modules in the co-expression network presented here may help to disentangle the effects of ageing *versus* the effects of a shift in behaviour, especially within solitary nests.

#### Genes highlighted both in differential expression analyses and the co-expression network represent particularly strong candidates for ageing research

The ten genes in solitary females convergently linked to ageing in both DEG and WGCN analyses are particularly good candidates to be further tested for their role in ageing processes. They were linked to immunity, venom production, gene expression and protein turnover, as well as fatty acid biosynthesis. Firstly, the increased expression of Toll-pathway inhibitors with age (*i. e*. the decreased activity of this immune pathway with age) suggests a negative relationship between age and immune defenses in solitary females. This may indicate a trade-off between reproduction and immunity in solitary females, as observed in many organisms and in line with the resource allocation model (van Noordwijk and de Jong 1986; Harshman and Zera 2007). Secondly, the higher expression of a gene related to venom production, *i.e*. venom acid phosphatase Acph-1, in young compared to old solitary females may indicate increased investment in defense early in life as females go through the nest-founding phase and must exit the nest to actively forage for their first brood. Again, following the resource allocation model, this investment may trade off with other life history traits in solitary females, such as somatic repair and longevity. Social dominant females may circumvent this trade-off, as in obligate eusocial species in which reproduction does not trade off against maintenance as a consequence of the abundant resources provided to the queen (Kramer et al. 2015). Further investigations would be necessary to confirm whether this is also the case in primitively eusocial species such as *E. viridissima*, but the pronounced transcriptomic signs of ageing in solitary compared to social individuals in this study seem to support this hypothesis. Thirdly, we found that a gene related to fatty acid biosynthesis (elongation of very long chain fatty acids) was linked to ageing in both of our analyses and was upregulated in young solitary females. Such genes have been associated with reduced lifespan in *C. elegans*, as higher expression of elongases is found in wild-type strains compared to long-lived mutants (Reis et al. 2011). Thus high expression levels of this gene in young solitary females may come at a cost for longevity. Finally, genes related to mRNA translation and protein turnover increased with age in solitary *E. viridissima* females. Although overall protein turnover rates are generally found to decrease with age in animals, leading to the accumulation of damaged proteins as organisms senesce (Ryazanov and Nefsky 2002), the dysregulation of certain transcription factors with age may lead to a “loss of silencing” and increased expression of particular genes (Frenk and Houseley 2018).

### No apparent role for juvenile hormone in *E. viridissima* ageing or social status

We report here the first data on JH titre levels in an orchid bee species. Strikingly, we found no significant difference in the haemolymph JH titres between young *vs* old and dominant *vs* subordinate females (Figure S17). So far, we cannot say much about the solitary *vs* social females due to the low sample number of solitary individuals (Figure S18).

On first sight, these findings apparently stand in contrast to the marked increase in JH levels in the honeybee workers, related with ageing and the transition from nursing to foraging tasks (Huang and Robinson 1996) and the equally strong positive correlation in JH levels with social status in the bumblebee *Bombus terrestris* (Bloch et al. 2000; Shpigler et al. 2014). In the stingless bee *Melipona quadrifasciata*, however, JH haemolymph titres in foragers are lower than those in workers (Cardoso-Junior et al. 2017). These evident discrepancies in the role of JH among representatives of the three social tribes of the monophyletic corbiculate bees, and now also a facultatively social euglossine, indicate that in corbiculate bees, which is the only clade within the bees that comprises members of all social complexities, there is no strong link between JH and reproduction, *i.e*. social status. Rather, each tribe appears to have moulded this ancient gonadotropic insect hormone (Santos, Humann, and Hartfelder 2019) according to its idiosyncratic mode of social biology. Such idiosyncrasies are not unique to the family Apidae (comprising the honeybees, bumblebees, orchid bees and stingless bees), but are also found in wasps. For instance, in the solitary progressively provisioning eumenine wasp *Synagris cornuta* there is also no correlation between JH levels and age or reproductive status (Kelstrup et al. 2018). Further, two sympatric species of the same genus of *Polistes* paper wasps showed a divergent patterns of JH titre with respect to age/task and social status (Kelstrup, Hartfelder, and Wossler 2015). Hence, the lack of a link between JH and age or social status in *E. viridissima* is not such a surprising result.

### Conclusions and future directions

The extensive changes in gene expression with age in solitary *Euglossa* females, which stand in stark contrast to the absence of gene expression changes with age in social *Euglossa* females, provide insight into the molecular pathways underlying ageing in solitary organisms, and how these may be modified in eusocial individuals. Our gene co-expression network further highlighted several hub genes correlated with age that were related to metabolic pathways, fatty acid biosynthesis and protein synthesis. The significant overlap of genes for which expression levels are associated with ageing across differential expression and co-expression network analyses revealed that a subset of genes related to immunity, venom production, protein production and fatty acid biosynthesis represent particularly relevant candidates for future ageing studies, both in solitary and social individuals. Ultimately, these results in a socially polymorphic species represent a tentative step in understanding the genetic mechanisms allowing the reversal of the fecundity/longevity trade-off which seemingly accompanies the evolutionary transition to eusociality (Carey 2001). Our study, based on expression differences inferred from transcriptome data, represents a useful starting point in the search for genetic markers and proximate mechanisms of ageing in the transition from solitary life to eusociality. A necessary next step in future studies would be to experimentally test the function of candidate genes identified here in order to eliminate possible confounding factors such as shifts in behaviour with age, for instance through knock-down experiments to corroborate or refute the role of these genes in relation to senescence.

Finally, in order to determine the relevance of these genes in the context of the fecundity/longevity trade-off, additional life history data are needed, such as information on the lifespan and reproductive output of individuals, to determine how these relate to expression of the candidate genes. Due to the invasive nature of RNA sampling from such small organisms, this is difficult to achieve. Emerging technologies may make this an accessible goal in the future, or studies in species for which larger sample sizes are available may allow parallel sampling of a subset of individuals alongside life history observations in the same or similar individuals.

## Acknowledgements

We thank Teresita Solís Sánchez for providing logistical support during fieldwork, particularly when marking bees. Particular thanks go to Constanze Wittkopp, Theresa Jörger-Hickfang, Atilla Çelikgil and Mert Ozle for their contributions in determining the cryptic species assignment of samples. We are grateful to all members of the So-Long research unit for helpful scientific discussions. This work was supported by the German Research Foundation (DFG) Research Unit FOR2281 (Sociality and the reversal of the fecundity/longevity trade-off) (components Pa 632/9, Ko 1895/20-1).

## Data Availability Statement

The data underlying this article are available in the National Centre for Biotechnology Information (NCBI) Sequence Read Archive (SRA), and will be accessible under Bioproject PRJNA636137, BioSample accession numbers SAMN15065573-SAMN15065604 upon publication of this manuscript. The genome annotation produced for *E. dilemma* (v2.2) will be available in the Dryad Digital Repository (doi: 10.5061/dryad.2547d7wnh) upon publication of this manuscript.

## References

Alonso-Alvarez C, Bertrand S, Devevey G, Prost J, Faivre B, Sorci G. 2004. Increased susceptibility to oxidative stress as a proximate cost of reproduction. Ecol Lett. 7: 363–368. https://doi.org/10.1111/j.1461-0248.2004.00594.x.

Altschul SF, Gish W, Miller W, Myers EW, Lipman DJ. 1990. Basic Local Alignment Search Tool. J Mol Biol. 215(3): 403–10.

Amdam GV, Aase ALTO, Seehuus SC, Fondrk MK, Norberg K, Hartfelder K. 2005. Social reversal of immunosenescence in honey bee workers. Exp Gerontol. 40(12): 939–47.

Andrade-Silva ACR, Nascimento FS. 2015. Reproductive regulation in an orchid bee: social context, fertility and chemical signalling. Anim Behav. 106: 43–49. https://doi.org/10.1016/j.anbehav.2015.05.004.

Andrews S. 2010. FastQC: A quality control tool for high throughput sequence data (version 0.11.4). https://www.bioinformatics.babraham.ac.uk/projects/fastqc/

Aumer D, Mumoki FN, Pirk CWW, Moritz RFA. 2018. The transcriptomic changes associated with the development of social parasitism in the honeybee Apis mellifera capensis. Sci Nat. 105: 22. https://doi.org/10.1007/s00114-018-1552-2.

Bahar R, Hartmann CH, Rodriguez KA, Denny AD, Busuttil RA, Dollé MET, Calder RB, Chisholm GB, Pollock BH, Klein CA, Vijg J. 2006. Increased cell-to-cell variation in gene expression in ageing mouse heart. Nature. 441(7096): 1011–14.

Barnes AI, Boone JM, Jacobson J, Partridge L, Chapman T. 2006. No extension of lifespan by ablation of germ line in Drosophila. Proc Biol Sci. 273(1589): 939–47.

Bates D, Mächler M, Bolker B, Walker S. 2015. Fitting linear mixed-effects models using lme4. J Stat Softw. https://doi.org/10.18637/jss.v067.i01.

Benjamini Y, Hochberg Y. 1995. Controlling the false discovery rate: a practical and powerful approach to multiple testing. J R Stat Soc Series B Stat Methodol. 57(1): 289–300. https://doi.org/10.1111/j.2517-6161.1995.tb02031.x.

Blagosklonny MV. 2010. Why men age faster but reproduce longer than women: mTOR and evolutionary perspectives. Aging. 2(5): 265–73.

Bloch G, Borst DW, Huang ZY, Robinson GE, Cnaani J, Hefetz A. 2000. Juvenile hormone titers, juvenile hormone biosynthesis, ovarian development and social environment in Bombus terrestris. J Insect Physiol. 46(1): 47–57.

Boff S, Saito CA, Alves-dos-Santos I. 2017. Multiple aggressions among nestmates lead to weak dominance hampering primitively eusocial behaviour in an orchid bee. Sociobiology. 64(2): 202–211. https://doi.org/10.13102/sociobiology.v64i2.1396.

Boff S, Forfert N, Paxton RJ, Montejo E, Quezada-Euan JJG. 2015. A behavioral guard caste in a primitively eusocial orchid bee, Euglossa viridissima, helps defend the nest against resin theft by conspecifics. Insectes Soc. 62: 247–249. https://doi.org/10.1007/s00040-015-0397-3.

Bolton A, Sumner S, Shreeves G, Casiraghi M, Field J. 2006. Colony genetic structure in a facultatively eusocial hover wasp. Behav Ecol. 17(6): 873–880. https://doi.org/10.1093/beheco/arl020.

Brand P, Hinojosa-Díaz IA, Ayala R, Daigle M, Yurrita Obiols CL, Eltz T, Ramírez SR. 2020. The evolution of sexual signaling is linked to odorant receptor tuning in perfume-collecting orchid bees. Nat Commun. 11 (1): 244.

Brand P, Saleh N, Pan H, Li C, Kapheim KM, Ramírez SRR. 2017. The nuclear and mitochondrial genomes of the facultatively eusocial orchid bee. G3 (Bethesda). 7 (9): 2891–98.

Bray NL, Pimentel H, Melsted P, Pachter L. 2016. Near-optimal probabilistic RNA-seq quantification. Nat Biotechnol. 34 (5): 525–27.

Cardoso-Júnior CAM, Silva RP, Borges NA, de Carvalho WJ, Walter SL, Simões ZLP, Bitondi MMG, Vieira CU, Bonetti AM, Hartfelder K. 2017. Methyl farnesoate epoxidase (mfe) gene expression and juvenile hormone titers in the life cycle of a highly eusocial stingless bee, Melipona scutellaris. J Insect Physiol. 101: 185–94.

Carey JR. 2001. Demographic mechanisms for the evolution of long life in social insects. Exp Gerontol. 36 (4-6): 713–22.

Charette JR, Samuels IS, Yu M, Stone L, Hicks W, Shi LY, Krebs MP, Naggert JK, Nishina PM, Peachey NS. 2016. A chemical mutagenesis screen identifies mouse models with ERG defects. Adv Exp Med Biol. 177–183. https://doi.org/10.1007/978-3-319-17121-0_24.

Cocom-Pech ME, de J. May-Itzá W, Medina Medina LA, Quezada-Euán JJG. 2008. Sociality in Euglossa (Euglossa) viridissima Friese (Hymenoptera, Apidae, Euglossini). Insectes Soc. 55(4): 428–433. https://doi.org/10.1007/s00040-008-1023-4.

Corona M, Velarde RA, Remolina S, Moran-Lauter A, Wang Y, Hughes KA, Robinson GE. 2007. Vitellogenin, juvenile hormone, insulin signaling, and queen honey bee longevity. Proc Natl Acad Sci U S A. 104 (17): 7128–33.

De Cecco M, Criscione SW, Peterson AL, Neretti N, Sedivy JM, Kreiling JA. 2013. Transposable elements become active and mobile in the genomes of aging mammalian somatic tissues. Aging. 5 (12): 867–83.

De Magalhães JP, Curado J, and Church GM. 2009. Meta-analysis of age-related gene expression profiles identifies common signatures of aging. Bioinformatics. 25 (7): 875–81.

De May-Itzá W, Medina Medina LA, Medina S, Paxton RJ, Quezada-Euán JJG. 2014. Seasonal nest characteristics of a facultatively social orchid bee, Euglossa viridissima, in the Yucatan Peninsula, Mexico. Insectes Soc. 61(2): 183–190. https://doi.org/10.1007/s00040-014-0342-x.

Dillin A, Crawford DK, Kenyon C. 2002. Timing requirements for insulin/IGF-1 signaling in C. Elegans. Science. 298(5594): 830–34.

Edward DA., Chapman T. 2011. Mechanisms underlying reproductive trade-offs: costs of reproduction. Mech Life Hist Evol. 137–152. https://doi.org/10.1093/acprof:oso/9780199568765.003.0011.

Elsner D, Meusemann K, Korb J. 2018. Longevity and transposon defense, the case of termite reproductives. Proc Natl Acad Sci U S A. 115(21): 5504–9.

Emms DM, Kelly S. 2015. OrthoFinder: solving fundamental biases in whole genome comparisons dramatically improves orthogroup inference accuracy. Genome Biol. 16: 157.

Fasken MB., Corbett AH, Stewart M. 2019. Structure-function relationships in the Nab2 polyadenosine-RNA binding Zn finger protein family. Prot Sci. 28(3): 513–23.

Festa-Bianchet M, King WJ. 1991. Effects of litter size and population dynamics on juvenile and maternal survival in Columbian ground squirrels. J Anim Ecol. 1077–1090. https://doi.org/10.2307/5432.

Flatt T. 2011. Survival costs of reproduction in Drosophila. Exp Gerontol. 46(5): 369–75.

Flatt T, Min KJ, D’Alterio C, Villa-Cuesta E, Cumbers J, Lehmann R, Jones DL, Tatar M. 2008. Drosophila germ-line modulation of insulin signaling and lifespan. Proc Natl Acad Sci U S A. 105(17): 6368–73.

Flatt T, Tu MP, Tatar M. 2005. Hormonal pleiotropy and the juvenile hormone regulation of Drosophila development and life history. Bioessays. 27 (10): 999–1010.

Freiria GA, Garófalo CA, Del Lama MA. 2017. The primitively social behavior of Euglossa cordata (Hymenoptera, Apidae, Euglossini): a view from the perspective of kin selection theory and models of reproductive skew. Apidologie. 48(4): 523–532. https://doi.org/10.1007/s13592-017-0496-4.

Frenk S, Houseley J. 2018. Gene expression hallmarks of cellular ageing. Biogerontology. 19(6): 547–66.

García-Velázquez L, Arias A. 2020. An update on the molecular pillars of aging. Clin Genet Genom Aging. 1–25. https://doi.org/10.1007/978-3-030-40955-5_1.

Harman, D. 1956. Aging: a theory based on free radical and radiation chemistry. J Gerontol. 11(3): 298–300.

Harman D. 1982. The free-radical theory of aging. Free Radic Biol. 275(3-6): 257–266. https://doi.org/10.1016/b978-0-12-566505-6.50015-6.

Harshman LG, Zera AJ. 2007. The cost of reproduction: the devil in the details. Trends Ecol Evol. 22(2): 80–86. https://doi.org/10.1016/j.tree.2006.10.008.

Hartfelder K, Engels W. 1998. Social insect polymorphism: hormonal regulation of plasticity in development and reproduction in the honeybee. Curr Top Dev Biol. 40: 45–77.

Hartfelder K, Bitondi MMG, Brent CS, Guidugli-Lazzarini KR, Simões ZLP, Stabentheiner A, Tanaka ED, Wang Y. 2013. Standard methods for physiology and biochemistry research in Apis mellifera. J Apic Res. 52(1): 1–48. https://doi.org/10.3896/ibra.1.52.1.06.

Huang ZY, Robinson GE. 1996. Regulation of honey bee division of labor by colony age demography. Behav Ecol Sociobiol. 39(3): 147–158. https://doi.org/10.1007/s002650050276.

Jiang H, Lei R, Ding SW, Zhu S. 2014. Skewer: a fast and accurate adapter trimmer for next-generation sequencing paired-end reads. BMC Bioinformatics. 15: 182.

Kapahi P, Chen D, Rogers AN, Katewa SD, Li PWL, Thomas EL, Kockel L. 2010. With TOR, less is more: a key role for the conserved nutrient-sensing TOR pathway in aging. Cell Metab. 11(6): 453–465. https://doi.org/10.1016/j.cmet.2010.05.001.

Kapheim KM. 2017. Nutritional, endocrine, and social influences on reproductive physiology at the origins of social behavior. Curr Opin Insect Sci. 22: 62–70.

Kapheim KM, Pan H, Li C, Salzberg SL, Puiu D, Magoc T, Robertson HM, Hudson ME, Venkat A, Fischman BJ, Hernandez A. 2015. Social evolution. Genomic signatures of evolutionary transitions from solitary to group living. Science. 348(6239): 1139–43.

Kapheim KM, Smith AR, Nonacs P, Wcislo WT, Wayne RK. 2013. Foundress polyphenism and the origins of eusociality in a facultatively eusocial sweat bee, Megalopta genalis (Halictidae). Behav Ecol Sociobiol. 67(2): 331–340. https://doi.org/10.1007/s00265-012-1453-x.

Kelstrup HC, Hartfelder K, Lopes TF, Wossler TC. 2018. The behavior and reproductive physiology of a solitary progressive provisioning vespid wasp: evidence for a solitary-cycle origin of reproductive castes. Am Nat. 191(2): E27–39.

Kelstrup HC, Hartfelder K, Wossler TC. 2015. Polistes smithii vs. Polistes dominula: the contrasting endocrinology and epicuticular signaling of sympatric paper wasps in the field. Behav Ecol Sociobiol. 69(12): 2043–2058. https://doi.org/10.1007/s00265-015-2015-9.

Kenyon C. 2010. A pathway that links reproductive status to lifespan in Caenorhabditis elegans. Ann N Y Acad Sci. 1204: 156–62.

Kramer BH, Schrempf A, Scheuerlein A, Heinze J. 2015. Ant colonies do not trade-off reproduction against maintenance. PLoS One. 10(9). https://doi.org/10.1371/journal.pone.0137969.

Kuhn JMM, Meusemann K, Korb J. 2019. Long live the queen, the king and the commoner? Transcript expression differences between old and young in the termite Cryptotermes secundus. PLoS One. 14(2). https://doi.org/10.1371/journal.pone.0210371.

Kuszewska K, Miler K, Rojek W, Woyciechowski M. 2017. Honeybee workers with higher reproductive potential live longer lives. Exp Gerontol. 98: 8–12.

Landis GN, Abdueva D, Skvortsov D, Yang J, Rabin BE, Carrick J, Tavaré S, Tower J. 2004. Similar gene expression patterns characterize aging and oxidative stress in Drosophila melanogaster. Proc Natl Acad Sci U S A. 101(20): 7663–68.

Langfelder P, Horvath SH. 2008. WGCNA: an R package for weighted correlation network analysis. BMC Bioinformatics. 9: 559.

Larsen PL. 1993. Aging and resistance to oxidative damage in Caenorhabditis elegans. Proc Natl Acad Sci U S A. 90(19): 8905–8909. https://doi.org/10.1073/pnas.90.19.8905.

Lin YJ, Seroude L, Benzer S. 1998. Extended life-span and stress resistance in the Drosophila mutant methuselah. Science. 282(5390): 943–46.

Li W, Prazak L, Chatterjee N, Grüninger S, Krug L, Theodorou D, Dubnau J. 2013. Activation of transposable elements during aging and neuronal decline in Drosophila. Nat Neurosci. 16(5): 529–31.

López-Otín C, Blasco MA, Partridge L, Serrano M, Kroemer G. 2013. The hallmarks of aging. Cell. 153(6): 1194–1217.

Love MI, Huber W, Anders S. 2014. Moderated estimation of fold change and dispersion for RNA-seq data with DESeq2. Genome Biol. 15(12): 550.

Lucas ER, Privman E, Keller L. 2016. Higher expression of somatic repair genes in long-lived ant queens than workers. Aging. 8(9): 1940–51.

Maynard-Smith J. 1958. The effects of temperature and of egg-laying on the longevity of Drosophila subobscura. Exp Biol. 35(4): 832–42.

Medawar PB. 1952. An unsolved problem of biology: inaugural lecture delivered at University College, London. H.K. Lewis and Company.

Monaghan P. 2014. Organismal stress, telomeres and life histories. J Exp Biol. 217 (1): 57–66.

NCBI nucleotide database. 2004. Bethesda (MD): National Library of Medicine (US), National Center for Biotechnology Information. [accessed 2020 01 07]. Available from: https://blast.ncbi.nlm.nih.gov/Blast.cgi

Negroni MA, Foitzik S, Feldmeyer B. 2019. Long-lived Temnothorax ant queens switch from investment in immunity to antioxidant production with age. Sci Rep. 9(1): 7270.

Page RE, Peng CY. 2001. Aging and development in social insects with emphasis on the honey bee, Apis mellifera L. Exp Gerontol. 36(4-6): 695–711.

Pérez VI, Bokov A, Van Remmen H, Mele J, Ran Q, Ikeno Y, Richardson A. 2009. Is the oxidative stress theory of aging dead? Biochim Biophys Acta. 1790(10): 1005–14.

Peters RS, Krogmann L, Mayer C, Donath A, Gunkel S, Meusemann K, Kozlov A, Podsiadlowski L, Petersen M, Lanfear R, Diez PA. 2017. Evolutionary history of the Hymenoptera. Curr Biol. 27(7): 1013–18.

Proshkina EN, Shaposhnikov MV, Sadritdinova AF, Kudryavtseva AV, Moskalev AA. 2015. Basic mechanisms of longevity: a case study of Drosophila pro-longevity genes. Ageing Res Rev. 24: 218–31.

R Core Team. 2019. R: A Language and Environment for Statistical Computing (version 3.6.1). https://www.R-project.org/.

Rawlings ND, Barrett AJ, Thomas PD, Huang X, Bateman A, Finn RD. 2018. The MEROPS database of proteolytic enzymes, their substrates and inhibitors in 2017 and a comparison with peptidases in the PANTHER database. Nucleic Acids Res. 44(D1): D343–D350. https://doi.org/10.1093/nar/gkx1134.

Redowicz MJ. 2007. Unconventional myosins in muscle. Eur J Cell Biol. 86(9): 549–558. https://doi.org/10.1016/j.ejcb.2007.05.007.

Reis RJS, Xu L, Lee H, Chae M, Thaden JJ, Bharill P, Tazearslan C, Siegel E, Alla R, Zimniak P, Ayyadevara S. 2011. Modulation of lipid biosynthesis vontributes to stress resistance and longevity of C. elegans mutants. Aging. 3(2): 125–47.

Rodrigues MA, Flatt T. 2016. Endocrine uncoupling of the trade-off between reproduction and somatic maintenance in eusocial insects. Curr Opin Insect Sci. 16: 1–8.

Ryazanov AG, Nefsky BS. 2002. Protein turnover plays a key role in aging. Mech Ageing Dev. 123(2-3): 207–13.

Saleh NW, Ramírez SRR. 2019. Sociality emerges from solitary behaviours and reproductive plasticity in the orchid bee Euglossa dilemma. Proc Biol Sci. 286(1906): 20190588.

Santos CG, Humann FC, Hartfelder K. 2019. Juvenile hormone signaling in insect oogenesis. Curr Opin Insect Sci. 31: 43–48.

Schrempf A, Giehr J, Röhrl R, Steigleder S, Heinze J. 2017. Royal Darwinian demons: enforced changes in reproductive efforts do not affect the life expectancy of ant queens. Am Nat. 189(4): 436–42.

Schrempf A, Heinze J, Cremer S. 2005. Sexual cooperation: mating increases longevity in ant queens. Curr Biol. 15(3): 267–70.

Seehuus SC, Norberg K, Gimsa U, Krekling T, Amdam GV. 2006. Reproductive protein protects functionally sterile honey bee workers from oxidative stress. Proc Natl Acad Sci U S A. 103(4): 962–67.

Séguret A, Bernadou A, Paxton RJ. 2016. Facultative social insects can provide insights into the reversal of the longevity/fecundity trade-off across the eusocial insects. Curr Opin Insect Sci. 16: 95–103.

Shell WA, Rehan SM. 2018. Behavioral and genetic mechanisms of social evolution: insights from incipiently and facultatively social bees. Apidologie. 49(1): 13–30. https://doi.org/10.1007/s13592-017-0527-1.

Shpigler H, Amsalem E, Huang ZY, Cohen M, Adam J. Siegel AJ, Hefetz A, Bloch G. 2014. Gonadotropic and physiological functions of juvenile hormone in bumblebee (Bombus terrestris) workers. PloS One. 9(6): e100650.

Soneson C, Love MI, Robinson MD. 2015. Differential analyses for RNA-seq: transcript-level estimates improve gene-level inferences. F1000Res. 4: 1521.

Southworth LK, Owen AB, Kim SK. 2009. Aging mice show a decreasing correlation of gene expression within genetic modules. PLoS Genet. 5(12): e1000776.

Tang H, Kambris Z, Lemaitre B, Hashimoto C. 2006. Two proteases defining a melanization cascade in the immune system of Drosophila. J Biol Chem. 281(38): 28097–104.

Todeschini AL, Teysset L, Delmarre V, Ronsseray S. 2010. The epigenetic trans-silencing effect in Drosophila involves maternally-transmitted small RNAs whose production depends on the piRNA pathway and HP1. PloS One. 5(6): e11032.

Toth AL, Sumner S, Jeanne RL. 2016. Patterns of longevity across a sociality gradient in vespid wasps. Curr Opin Insect Sci. 16: 28–35.

Van Noordwijk AJ, de Jong G. 1986. Acquisition and allocation of resources: their influence on variation in life history tactics. Am Nat. 128(1): 137–142. https://doi.org/10.1086/284547.

Vu VQ. 2011. Ggbiplot: A ggplot2 Based Biplot (R package version 0.55). http://github.com/vqv/ggbiplot.

Wang Y, Salmon AB, Harshman LG. 2001. A cost of reproduction: oxidative stress susceptibility is associated with increased egg production in Drosophila melanogaster. Exp Gerontol. 36(8): 1349–1359. https://doi.org/10.1016/s0531-5565(01)00095-x.

Williams GC. 1957. Pleiotropy, natural selection, and the evolution of senescence. Evolution 11:398–411.

Zangar RC, Davydov DR, Verma S. 2004. Mechanisms that regulate production of reactive oxygen species by cytochrome P450. Toxicol Appl Pharmacol. 199(3): 316–31.

Zhang B, Horvath S. 2005. A general framework for weighted gene co-expression network analysis. Stat Appl Genet Mol Biol. 4: Article17.

Zhang Q, Skepper JN, Yang F, Davies JD, Hegyi L, Roberts RG, Weissberg PL, Ellis JA, Shanahan CM. 2001. Nesprins: a novel family of spectrin-repeat-containing proteins that localize to the nuclear membrane in multiple tissues. J Cell Science. 114: 4485–98.

Zhu M, Zhang W, Liu F, Chen X, Li H, Xu B. 2016. Characterization of an Apis cerana cerana cytochrome P450 gene (AccCYP336A1) and its roles in oxidative stresses responses. Gene. 584(2): 120–28.

