## Supplemental Files S1, S3-S5, S7, S8, S12, S16-S18 for "Transcriptomic signatures of ageing vary in solitary and social forms of an orchid bee"

### Supplementary material

Table S1. Behaviours (a) and positions in the nest (b) recorded in order to determine the phenotype (dominant *vs* subordinate) of females in social nests. The phenotype was determined after observing the nest every five minutes over multiple half-hour periods until roles (dominant *vs* subordinate) could be unambiguously assigned.

a)

| Behaviour | Associated phenotype |
|---|---|
| foraging | subordinate |
| guarding entrance | subordinate (guard) |
| waving antennae towards other female | dominant |
| touching other female | dominant |
| launching herself at other female | dominant |
| "kicking" other female | dominant |
| egg-laying | dominant |

b)

| Location | Associated phenotype |
|---|---|
| on brood cells | dominant |
| at nest entrance | subordinate (guard) |
| on nest floor | subordinate |

Table S3. Species assignment of samples to *E. dilemma* or *E. viridissima*. The assignment was done based on Sanger sequencing of the *or41* gene (Brand *et al.* 2020). All individuals in our study were homozygous across all six single nucleotide polymorphisms known to differentiate the two species in males, making the assignment of our diploid female samples unambiguous.

| sample ID | species |
|---|---|
| eug1 | *E. viridissima* |
| eug2 | *E. dilemma* |
| eug3 | *E. dilemma* |
| eug4 | *E. dilemma* |
| eug5 | *E. dilemma* |
| eug6 | *E. viridissima* |
| eug7 | *E. viridissima* |
| eug8 | *E. viridissima* |
| eug9 | *E. viridissima* |
| eug10 | *E. viridissima* |
| eug11 | *E. viridissima* |
| eug12 | *E. viridissima* |
| eug13 | *E. viridissima* |
| eug14 | *E. viridissima* |
| eug15 | *E. dilemma* |
| eug16 | *E. viridissima* |
| eug17 | *E. viridissima* |
| eug18 | *E. viridissima* |
| eug19 | *E. viridissima* |
| eug20 | *E. viridissima* |
| eug21 | *E. viridissima* |
| eug22 | *E. viridissima* |
| eug23 | *E. viridissima* |
| eug24 | *E. viridissima* |
| eug25 | *E. viridissima* |
| eug26 | *E. viridissima* |
| eug27 | *E. viridissima* |
| eug28 | *E. viridissima* |
| eug29 | *E. viridissima* |
| eug30 | *E. viridissima* |
| eug31 | *E. viridissima* |
| eug32 | *E. viridissima* |
| eug33 | *E. viridissima* |
| eug34 | *E. dilemma* |

Brand P, Hinojosa-Díaz IA, Ayala R, Daigle M, Obiols CLY, Eltz T, Ramirez SR. 2020. The evolution of sexual signaling is linked to odorant receptor tuning in perfume-collecting orchid bees. Nat Commun. 11. doi:10.1038/s41467-019-14162-6

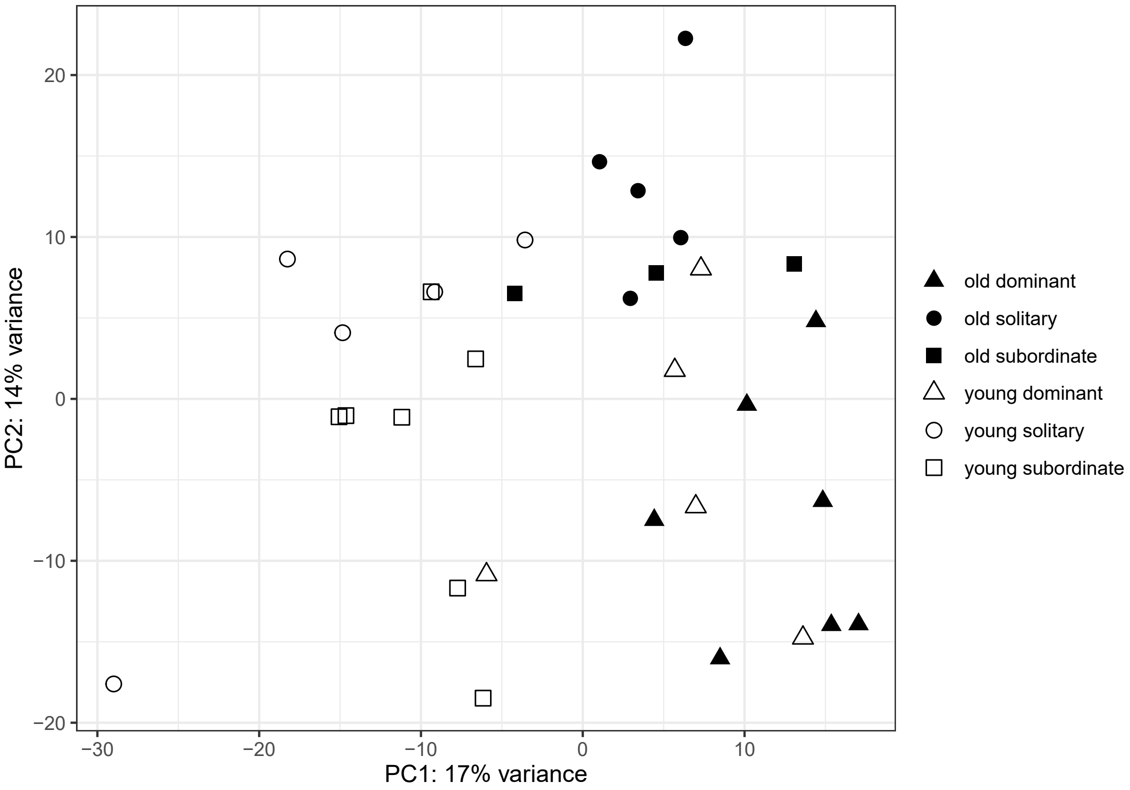


Fig. S4. Principal component analysis (PCA) of variance-stabilised RNA read counts for young and old females, from solitary and social nests. All individuals sequenced, including *E. dilemma* individuals and the one outlier (young solitary, bottom left), are represented in this figure. Each point represents the expression profile across all genes for one individual. Axis labels indicate the amount of variance in gene expression explained by the first two principal components (PC1 and PC2).

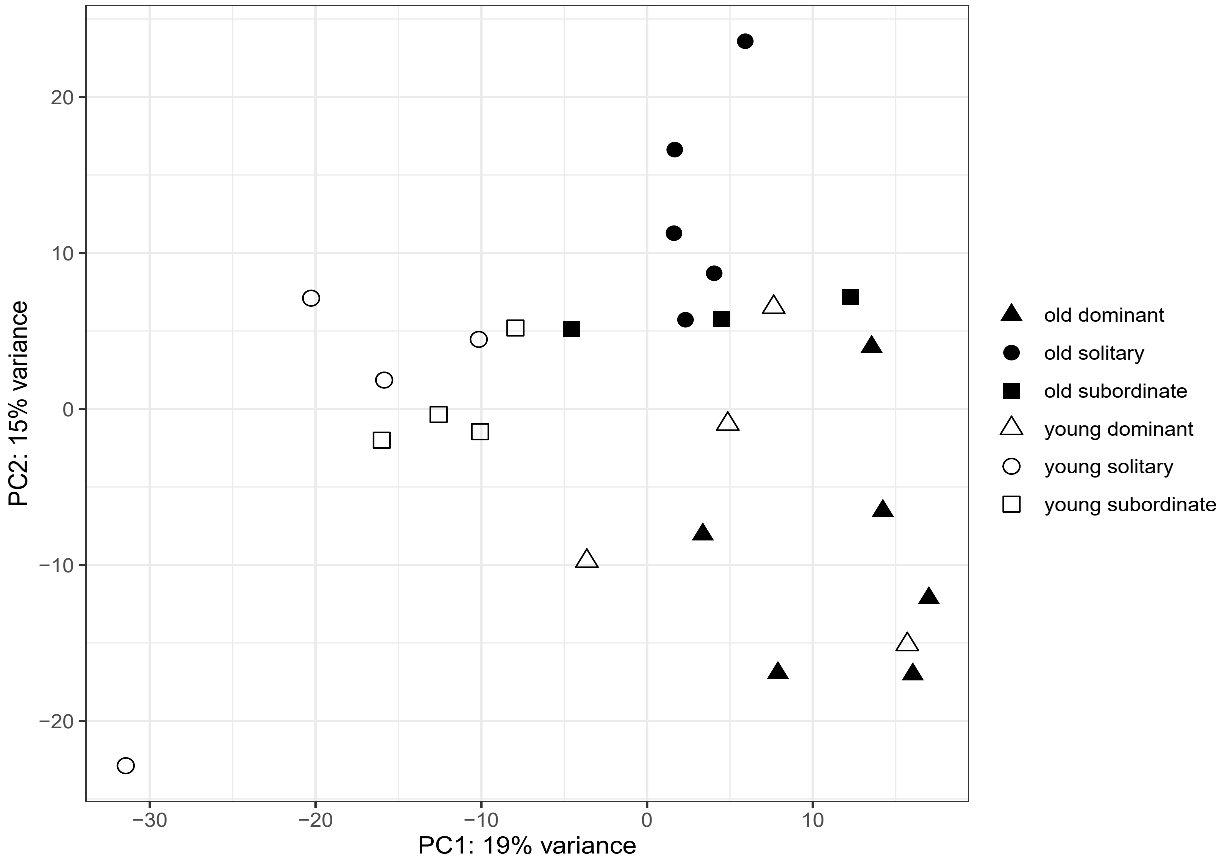


Fig. S5. Principal component analysis (PCA) of variance-stabilised RNA read counts for young and old females, from solitary and social nests. All E. viridissima individuals are represented here, including the one outlier (young solitary, bottom left). Each point represents the expression profile across all genes for one individual. Axis labels indicate the amount of variance in gene expression explained by the first two principal components (PC1 and PC2).

Table S7. Number of significantly differentially expressed genes (DEGs, adjusted $p < 0.05$) for each pairwise comparison across age groups and social types. The analysis including all individuals (*E. viridissima* including the one outlier, and *E. dilemma* individuals) yielded similar numbers to the analysis only including "core individuals" (excluding *E. dilemma* and the one outlier, sample "eug6").

| pairwise comparison | number of significant DEGs all individuals | number of significant DEGs core individuals |
|---|---|---|
| solitary young *vs* old | 792 | 940 |
| dominant young *vs* old | 5 | 14 |
| subordinate young *vs* old | 8 | 7 |
| young subordinate *vs* solitary | 13 | 38 |
| young subordinate *vs* dominant | 218 | 212 |
| old subordinate *vs* solitary | 32 | 53 |
| old subordinate *vs* dominant | 19 | 38 |

File S8. Detailed description of the pipeline for transcriptome assembly and *E. dilemma* genome annotation Edil_v2.2.

The transcriptomic analyses are based on the previously published draft genome sequence assembly for *E. dilemma* (GCA_002201625.1, (Brand et al. 2017)). Repeats were softmasked (35.30% of the total genome assembly length) using bedtools (v2.27.1) based on repeat annotations from Tandem Repeats Finder (v4.09, (Benson 1999)) and RepeatMasker (Smit, AFA, Hubley, R & Green, P. 2013-2020). To improve the previous annotation of this genome, which was based solely on gene predictions and homology to *Apis mellifera* proteins (Brand et al. 2017), we used Funannotate (v1.5.1, github.com/nextgenusfs/funannotate) with the previous gene annotation (edil.1.0.annotations.gff, (Brand et al. 2017)), novel experimental evidence derived from RNA-seq data (generated in this study: RNA-seq paired-end reads (euglossa.R[1,2].fastq.gz) aligned to the *E. dilemma* genome with HiSat2 (v2.1.0, (Kim, Langmead, and Salzberg 2015)) in conjunction with processing with samtools (v1.8, (Li et al. 2009)) and sambamba (v0.7.1, (Tarasov et al. 2015)) (euglossa.all.hisat2.bam), transcriptome assembly with binpacker (v1.1, (Liu et al. 2016)) from 4 samples (binpacker.transcriptome.fa), genome-guided transcriptome assembly from one sample (eug33, stringtie2.transcriptome.gtf) with stringtie2 (v1.3.3b, (M. Pertea et al. 2016))), a RNA-seq data-based Trinity (v1.5.1, (Grabherr et al. 2011)) transcriptome assembly of the closely related *E. dilemma* (euglossa.dilemma.trinity.transcriptome.fa, P. Brand, pers. comm., data published in Brand et al. 2020), transcriptomes of the closely related orchid bee *Eufrisea mexicana* (GCF_001483705.1 ASM148370v1_rna.fna). Further, protein sequences from 5 related bee species (*Bombus impatiens*: GCF_000188095.2 BIMP_2.1_protein.faa; *B. terrestris*: GCF_000214255.1 Bter_1.0_protein.faa; *Apis mellifera*: GCF_003254395.2 Amel_HAv3.1_protein.faa; *Melipona quadrifasciata*: GCA_001276565.1 ASM127656v1_protein.faa; *Eufrisea mexicana*: GCF_001483705.1 ASM148370v1_protein.faa, (Wallberg et al. 2019; Kapheim et al. 2015)), and Uniprot (sprot) was used for homology-based evidence. Additionally, genes were predicted from the *E. dilemma* genome using SNAP (v2006-07-28, (Korf 2004)) based on SNAP's *Apis mellifera* HMM dataset (snap.predicted.gff), Genemark-ET (--max_intron 3000, (Borodovsky and Lomsadze 2011)), Augustus (v3.3, (Hoff and Stanke 2019)) (--genemodel=partial --strand=both --maxDNAPieceSize=200000 --protein=on --progress=true --uniqueGeneId=true --stopCodonExcludedFromCDS=False --gff3=on --species=Euglossa_dilemma) based on BUSCO3 (v3.0.2, (Simão et al. 2015)) training (-m geno --long --species bombus_impatiens1, database: Hymenoptera odb9), and the PASA pipeline (v2.3.3, (B. J. Haas et al. 2008), --MAX_INTRON_LENGTH 10000 --ALIGNERS blat,gmap --stringent_alignment_overlap 50 --trans_gtf stringtie2.transcriptome.gtf -t euglossa.dilemma.trinity.transcriptome.fa).

In brief, PASA (v2.3.3, (B. J. Haas et al. 2008)) was used to train gene predictions (funannotate train --max_intronlen 5000 --pasa_alignment_overlap 50.0 --coverage 50 --stranded no, euglossa.dilemma.trinity.transcriptome.fa, euglossa.R[1,2].fastq.gz). With this training set, Funannotate (v1.5.1) prediction was performed (funannotate predict --repeat_filter overlap blast --busco_seed_species Euglossa_dilemma --rna_bam euglossa.all.hisat2.bam --ploidy 1 --soft_mask 5000 --repeats2evm --stringtie stringtie2.transcriptome.gtf --other_gff snap.predicted.gff --other_gff edil.1.0.annotations.gff:8 --protein_evidence uniprot/5.Bee.proteomes.faa --transcript_evidence euglossa.dilemma.trinity.transcriptome.fa binpacker.transcriptome.fa edil.1.0.annotations.fa Eufriesea.mexicana.RNA.fna). In this step transcriptomes and proteomes are aligned to the *E. dilemma* genome with minimap2 (v2.17-r954, (Li 2018)) and proteins aligned with Diamond (v0.8, (Buchfink, Xie, and Huson 2015)) and exonerate (v2.4.0, (Slater and Birney 2005)), gene predictions are generated based on the genome by Genemark-ES ((Borodovsky and Lomsadze 2011)) and

Augustus (v.3.3, (Hoff and Stanke 2019)), based on RNA-seq alignments by CodingQuarry (v2.0, (Testa et al. 2015)), and then integrated other input annotations in Evidence Modeler (v.1.1.1, (Testa et al. 2015; B. J. Haas et al. 2008)) to a total of 23,819 gene models. Too short, gap-spanning or repeat-overlapping gene models were removed so that 23,093 gene models remained. Additionally, 185 tRNA gene models were found with tRNAscan.

Gene models are then updated with PASA and transdecoder (v5.5.0, Haas and Papanicolaou (B. J. A. P. Haas 2019)) (funannotate update --max_intronlen 6000 --pasa_alignment_overlap 30.0 --coverage 30), using empirical evidence (RNA-Seq reads: euglossa.R[1,2].fastq.gz, Trinity transcriptome assembly: euglossa.dilemma.trinity.transcriptome.fa). For 23,093 gene models and PASA aligned transcripts, Kallisto (v0.46.2) TPM value were used to update gene models and to determine which PASA gene models to select at each locus. Two problematic gene models were fixed (funannotate fix).

Then genes were functionally annotated (funannotate iprscan --method local --cpus 110 --num 1000 --out funannotate.train1/interproscan.output.xml; funannotate annotate --busco_db hymenoptera, --iprscan funannotate.train1/interproscan.output.xml), including HMMer (v2.3.2, v3.1b2, (Mistry et al. 2013)) search of PFAM version 32.0 ((El-Gebali et al. 2019)), Diamond (v0.8, (Buchfink, Xie, and Huson 2015)) blastp search of UniProt DB version 2018_11, EggNog Annotations (eggnog mapper v1.0.3 (Huerta-Cepas et al. 2017), eggnog_4.5/hmmdb databases: arthopoda, insecta, hymanoptera, drosophila, database created with "download_eggnog_data.py euk artNOG inNOG meNOG droNOG hymNOG" and "diamond.0.8 getseq --db eggnog_proteins_old.dmnd | diamond makedb --db eggnog_proteins.dmnd"), Diamond (v0.8) blastp search of MEROPS version 12.0 (Rawlings et al. 2018), annotating CAZYmes using HMMer search of dbCAN version 7.0 (Huang et al. 2018), annotating proteins with BUSCO hymenoptera models (v3.0.2, Hymenoptera odb9, (Simão et al. 2015)), predicting secreted proteins with SignalP (4.1, (Nielsen 2017)), InterProScan5 annotations (v5.33-72.0, (Jones et al. 2014)).

The resulting set of 23,340 protein coding genes and 125 tRNA genes were processed, checked for quality and compared to the previous *E. dilemma* annotation (v1.0), using BUSCO3 (v3.0.2, (Simão et al. 2015)), transvestigator (v0.1-alpha), gff3_gene_prediction_file_validator.pl (Evidence Modeler, EvmUtils) gff3/gff3validator (genometools, v1.5.10, (Gremme, Steinbiss, and Kurtz 2013)), gffcompare/gffread/GFFcleaner (GFF Utilities, v0.10.4, (G. Pertea and Pertea 2020)), gff3_QC (gff3 toolkit, v1.4.4), transeq (EMBOSS, v6.6.0, (Rice, Longden, and Bleasby 2000)), xtractore (AEGeAn toolkit, v0.16.0, (Standage 2015)) and gene validator (v2.1.9, (Drăgan et al. 2016)). This comparison showed that a number of gene models were filtered, mainly due to stop codons and gap spanning. Since some of these models are part of the conserved set of single copy orthologs (BUSCO), we extracted non-duplicated *E. dilemma* v1.0 annotations and rerun funannotate with these, and additionally with PASA pipeline predictions, BUSCO-trained Augustus predictions as additional input.

Again, PASA was used to train gene predictions (see above). With this training set, Funannotate prediction was performed (see above, --busco_seed_species bombus_impatiens1 --busco_db hymenoptera; modified funannotate-predict.py: --keep_no_stops, Busco weight = 5 (from 2), weights HiQ= 8 (from 5)). In this step transcriptomes and proteomes are aligned to the *E. dilemma* genome with minimap2 (136,041 alignments) and proteins (n=645,086) aligned with diamond (816,608 alignments) and exonerate (31,628 alignments), gene predictions are generated based on the genome by Genemark-ES (59,675 gene models) and Augustus (14,835 gene models, of which 6,332 are high quality predictions, *i.e.* > 90% exon evidence), based on RNA-seq alignments by CodingQuarry

(21,826 gene models), based on PASA predictions (9,911 gene models), and then integrated other input annotations (15,904 gene models) (total of 122,151 gene models) in Evidence Modeler to a total of 24,526 gene models. Too short, gap-spanning or repeat-overlapping gene models were removed (72,25,634 respectively, total 723) so that 23,803 gene models remained. Additionally, 185 tRNA gene models were found, of which 126 were valid (non-overlapping, tRNAscan SE v2.0.0, (Chan and Lowe 2019)).

Gene models were then updated with PASA and transdecoder (see above). With the selected 24,007 transcripts derived from 23,892 protein coding loci, a total 24,015 loci were written (24,015 loci, 24,128 transcripts, 167 new gene models, 18,827 unchanged models, 4,982 models with updated UTRs, 35 exons changed, 4 exons/CDS changed). One problematic gene model was fixed (FUN_012919 Feature begins or ends in a gap starting at 468,722).

Then genes were functionally annotated (see above), including HMMer search of PFAM version 32.0 (14,623 annotations), Diamond blastp search of UniProt DB version 2018_11 (345 valid gene/product annotations from 11,355 total), EggNog Annotations (22,708 COG/EggNog annotations, combined with the UniProt results: 1,758 gene name and product description annotations added), Diamond blastp search of MEROPS version 12.0 (433 annotations), annotating CAZYmes using HMMer search of dbCAN version 7.0 (321 annotations), annotating proteins with BUSCO hymenoptera models (3,627 annotations), predicting secreted proteins with SignalP (1,261 secretome and 0 transmembrane annotations), InterProScan5 annotations (99,658 valid annotations, 0 duplications).

The resulting set of gene annotations was processed and checked for quality (see above). Again, the comparison to the *E. dilemma* v1.0 annotation revealed a proportion of 1.0 annotations not present in the Funannotate annotations. Of these, 13 non-redundant transcripts from a total of 72 duplicated transcripts, 268 transcripts for which Proteins have internal stop codons, and 16 transcripts with Ns in a feature were added to the Funannotate results, totalling 24,991 gene models (deposited in Dryad, doi:10.5061/dryad.2547d7wnh). BUSCO3 found 3,560 out of 4,415 (80.6%) conserved single copy Hymenoptera orthologs to be complete, of which 89 (2.0%) were duplicated, 428 (9.7%) were fragmented, and 427 (9.7%) were missing.

### References

Benson G. 1999. Tandem Repeats Finder: a program to analyze DNA sequences. Nucleic Acids Res. 27(2): 573-580. https://doi.org/10.1093/nar/27.2.573.

Borodovsky M, Lomsadze A. 2011. Eukaryotic gene prediction using GeneMark.hmm-E and GeneMark-ES. Curr Protoc Bioinformatics. 35(1): 4.6.1-4.6.10. https://doi.org/10.1002/0471250953.bi0406s35.

Brand P, Saleh N, Pan H, Li C, Kapheim KM, Ramírez SR. 2017. The nuclear and mitochondrial genomes of the facultatively eusocial orchid bee *Euglossa dilemma*. G3 (Bethesda). 7(9): 2891-2898. https://doi.org/10.1101/123687.

Buchfink B, Xie C, Huson DH. 2015. Fast and sensitive protein alignment using DIAMOND. Nat Methods. 12(1): 59–60.

Chan PP, Lowe TM. 2019. tRNAscan-SE: searching for tRNA genes in genomic sequences. Methods Mol Biol. 1962: 1–14.

Drăgan MA, Moghul I, Priyam A, Bustos C, Wurm Y. 2016. GeneValidator: identify problems with protein-coding gene predictions. Bioinformatics. 32(10): 1559–61.

El-Gebali S, Mistry J, Bateman A, Eddy SR, Luciani A, Potter SC, Qureshi M, Richardson LJ, Salazar GA, Smart A, Sonnhammer ELL. 2019. The Pfam protein families database in 2019.

Nucleic Acids Res. 47(D1): D427–32.
Grabherr MG, Haas BJ, Yassour M, Levin JZ, Thompson DA, Amit I, Adiconis X, Fan L, Raychowdhury R, Zeng Q, Chen Z, Mauceli E, Hacohen N, Gnirke A, Rhind N, di Palma F, Birren BW, Nusbaum C, Lindblad-Toh K, Friedman N, Regev A. 2011. Full-length transcriptome assembly from RNA-Seq data without a reference genome. Nat Biotechnol. 29(7):644-52. doi: 10.1038/nbt.1883.
Gremme G, Steinbiss S, Kurtz S. 2013. GenomeTools: a comprehensive software library for efficient processing of structured genome annotations. IEEE/ACM Trans Comput Biol Bioinform. 10(3): 645–56.
Haas BJ, Papanicolaou A. 2019. *TransDecoder* (version v5.5.0). https://github.com/TransDecoder/TransDecoder/wiki.
Haas BJ, Salzberg SL, Zhu W, Pertea M, Allen JE, Orvis J, White O, Buell CR, Wortman JR. 2008. Automated eukaryotic gene structure annotation using EVidenceModeler and the program to assemble spliced alignments. Genome Biol. 9(1): R7.
Hoff KJ, Stanke M. 2019. Predicting genes in single genomes with AUGUSTUS. Curr Protoc Bioinformatics. 65(1): e57.
Huang L, Zhang H, Wu P, Entwistle S, Li X, Yohe T, Yi H, Yang Z, Yin Y. 2018. dbCAN-Seq: A database of carbohydrate-cctive enzyme (CAZyme) sequence and annotation. Nucleic Acids Res. 46(D1): D516–21.
Huerta-Cepas J, Forslund K, Coelho LP, Szklarczyk D, Jensen LJ, von Mering C, Bork P. 2017. Fast genome-wide functional annotation through orthology assignment by eggNOG-Mapper. Mol Biol Evol. 34(8): 2115–22.
Jones P, Binns D, Chang HY, Fraser M, Li W, McAnulla C, McWilliam H, Maslen J, Mitchell A, Nuka G, Pesseat S. 2014. InterProScan 5: genome-scale protein function classification. Bioinformatics. 30(9): 1236–40.
Kapheim KM, Pan H, Li C, Salzberg SL, Puiu D, Magoc T, Robertson HM, Hudson ME, Venkat A, Fischman BJ, Hernandez A. 2015. Social evolution. Genomic signatures of evolutionary transitions from solitary to group living. Science. 348(6239): 1139–43.
Kim D, Langmead B, Salzberg SL. 2015. HISAT: A fast spliced aligner with low memory requirements. Nat Methods. 12(4): 357–60.
Korf I. 2004. Gene finding in novel genomes. BMC Bioinformatics. 5: 59.
Li H. 2018. Minimap2: pairwise alignment for nucleotide sequences. Bioinformatics. 34(18): 3094-3100. https://doi.org/10.1093/bioinformatics/bty191.
Li H, Handsaker B, Wysoker A, Fennell T, Ruan J, Homer N, Marth G, Abecasis G, Durbin R, 1000 Genome Project Data Processing Subgroup. 2009. The sequence alignment/map format and SAMtools. Bioinformatics. 25(16): 2078–79.
Liu J, Li G, Chang Z, Yu T, Liu B, McMullen R, Chen P, Huang X. 2016. BinPacker: packing-based de novo transcriptome assembly from RNA-seq data. PLoS Comp Biol. 12(2): e1004772.
Mistry J, Finn RD, Eddy SR, Bateman A, Punta M. 2013. Challenges in homology search: HMMER3 and convergent evolution of coiled-coil regions. Nucleic Acids Res. 41(12): e121.
Nielsen H. 2017. Predicting secretory proteins with SignalP. Methods Mol Biol. 1611: 59–73.
Pertea G, Pertea M. 2020. GFF utilities: GffRead and GffCompare. F1000Res. 9(304): 304. https://doi.org/10.12688/f1000research.23297.1.
Pertea M, Kim D, Pertea GM, Leek JT, Salzberg SL. 2016. Transcript-level expression analysis of RNA-seq experiments with HISAT, StringTie and Ballgown. Nat Protoc. 11(9): 1650–67.
Rawlings ND, Barrett AJ, Thomas PD, Huang X, Bateman A, Finn RD. 2018. The MEROPS database of proteolytic enzymes, their substrates and inhibitors in 2017 and a comparison with peptidases in the PANTHER database. Nucleic Acids Res. 46(D1): D624-D632. https://doi.org/10.1093/nar/gkx1134.
Rice P, Longden I, Bleasby A. 2000. EMBOSS: the European molecular biology open software suite. Trends Genet. 16(6): 276–77.
Simão FA, Waterhouse RM, Ioannidis P, Kriventseva EV, Zdobnov EM. 2015. BUSCO: assessing genome assembly and annotation completeness with single-copy orthologs. Bioinformatics. 31(19): 3210–12.
Slater GSC, Birney E. 2005. Automated generation of heuristics for biological sequence comparison.

BMC Bioinformatics. 6: 31.
Smit AFA, Hubley R, Green P. 2013-2020. RepeatMasker (version 4.0). http://www.repeatmasker.org.
Standage DS. 2015. AEGeAn: an integrated toolkit for analysis and evaluation of annotated genomes (version v0.16.0). http://standage.github.io/AEGeAn.
Tarasov A, Vilella AJ, Cuppen E, Nijman IJ, Prins P. 2015. Sambamba: fast processing of NGS alignment formats. Bioinformatics. 31(12): 2032–34.
Testa AC, Hane JK, Ellwood SR, Oliver RP. 2015. CodingQuarry: highly accurate Hidden Markov Model gene prediction in fungal genomes using RNA-seq transcripts. BMC Genomics. 16: 170.
Wallberg A, Bunikis I, Pettersson OV, Mosbech MB, Childers AK, Evans JD, Mikheyev AS, Robertson HM, Robinson GE, Webster MT. 2019. A hybrid de novo genome assembly of the honeybee, *Apis mellifera*, with chromosome-length scaffolds. BMC Genomics. 20(1): 275.

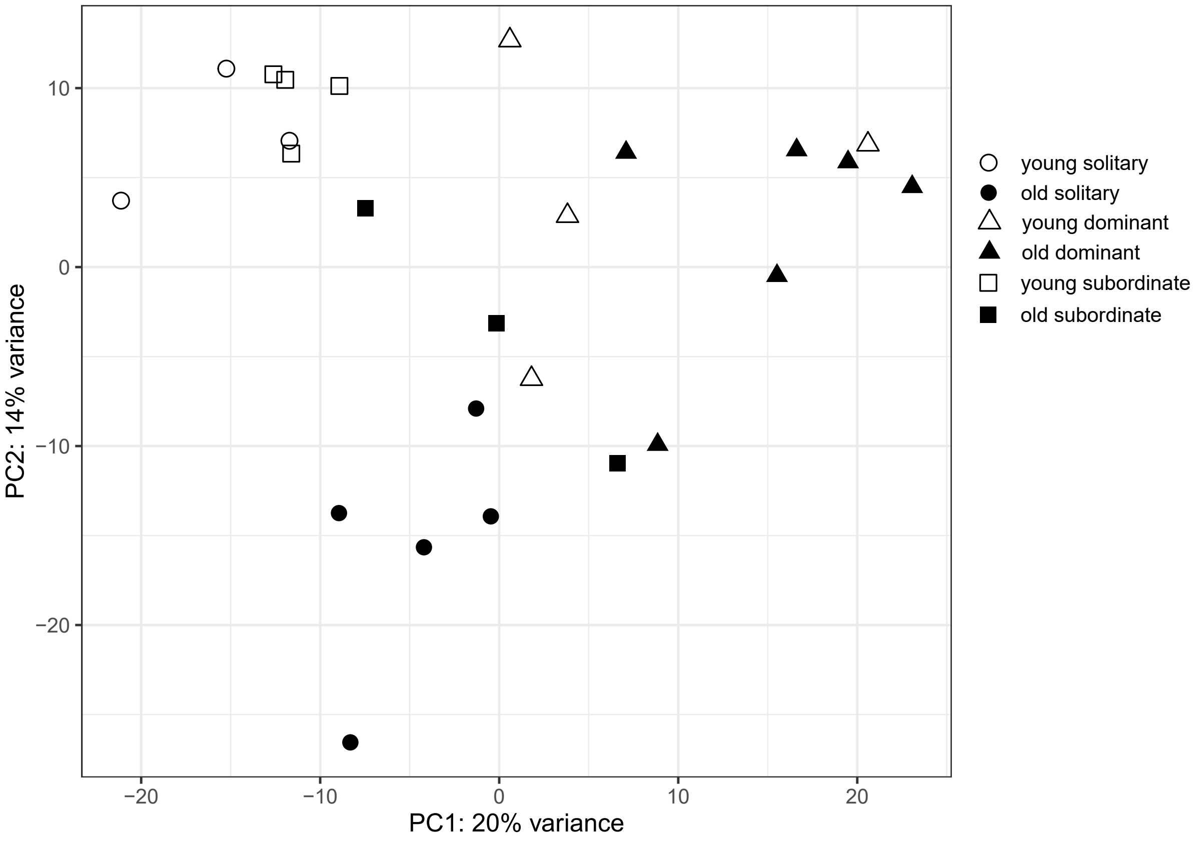


Fig. S12. Principal component analysis (PCA) of variance-stabilised RNA read counts for young and old females, from solitary and social nests. Each point represents the expression profile across all genes for one individual. Axis labels indicate the amount of variance in gene expression explained by the first two principal components (PC1 and PC2).

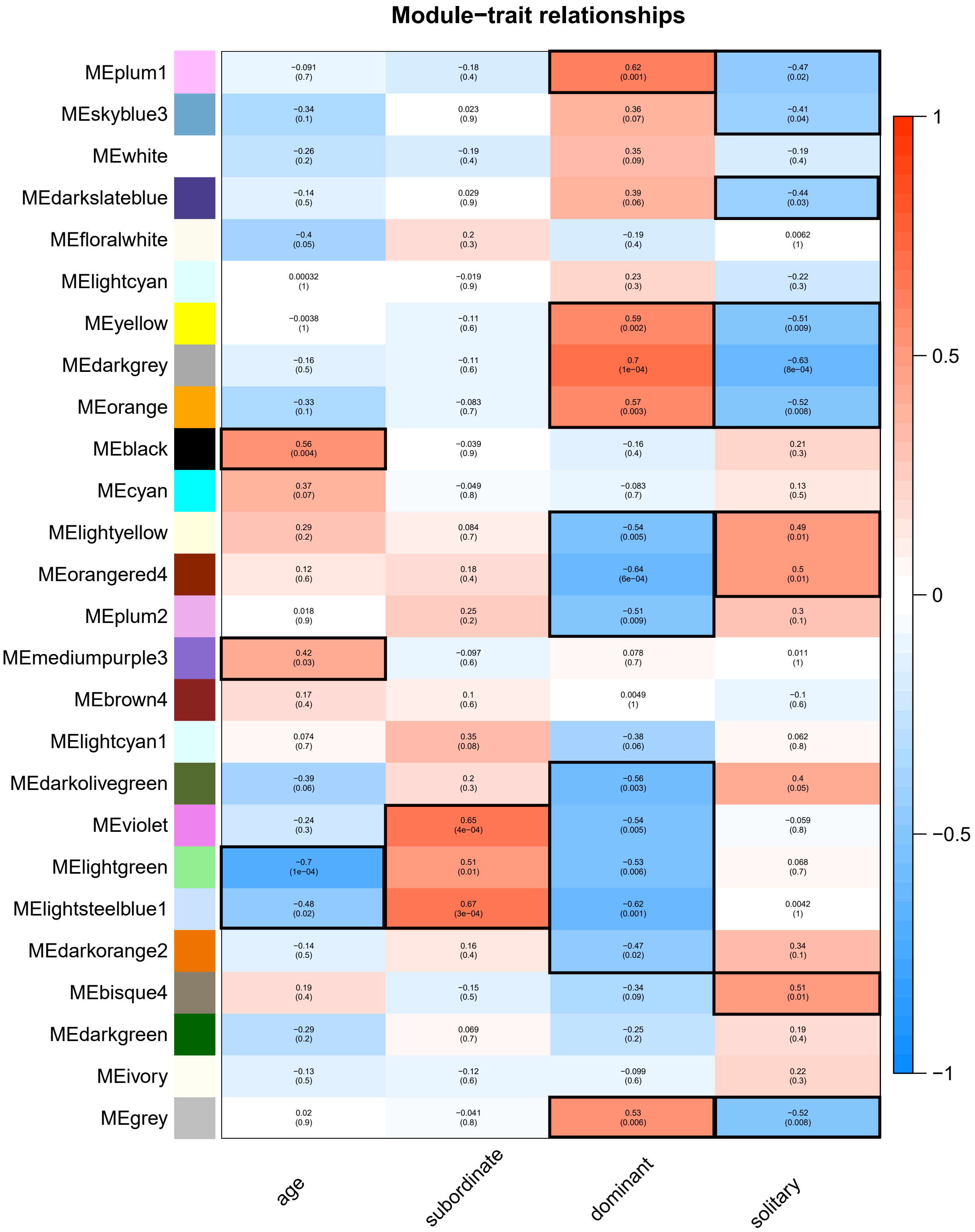


Fig. S16. Heatmap representing the correlations between module eigengenes and external traits of interest: age and social status (subordinate, dominant or solitary). Numbers shown in each square represent the correlation coefficient, with the associated p-value in brackets. Red squares indicate a positive correlation between the module eigengene and the trait; blue squares indicate a negative correlation. Modules with a p-value < 0.05 for a particular trait (highlighted by a black outline) were considered to be significantly correlated with that trait.

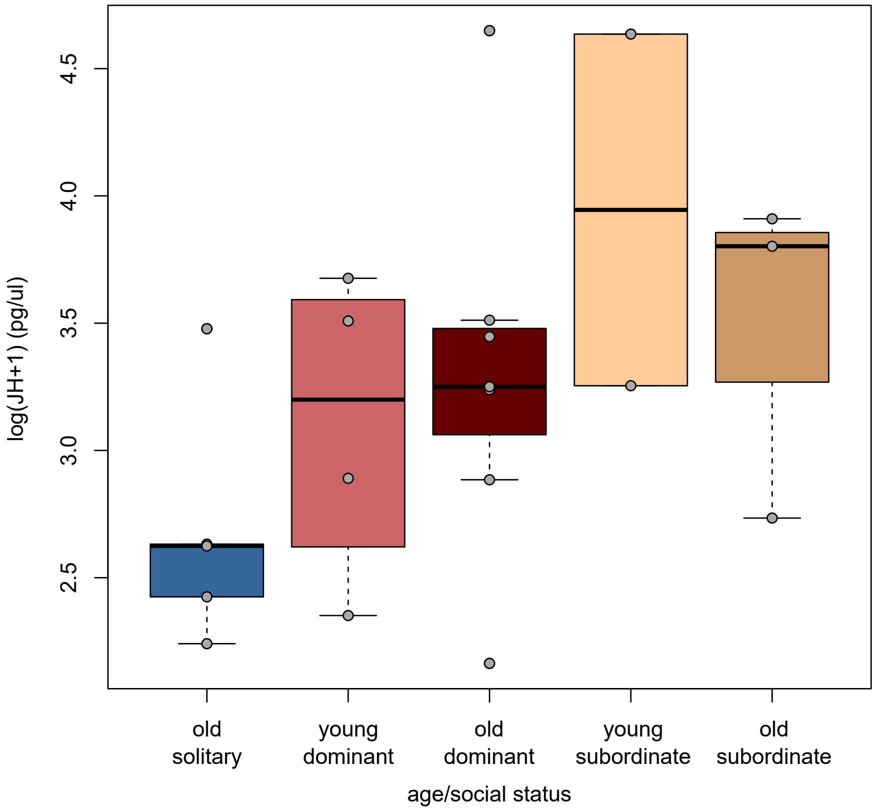


Fig. S17. Boxplot (median ± interquartile range) of log-transformed juvenile hormone titres (pg/µl) for females of different age and social status (no juvenile hormone measurements could be made for young solitary females). *E. dilemma* individuals were excluded from this analysis. Points indicate individual titres. The differences observed between groups were not statistically significant (LMM, age: $\chi^2(1,21) = 0.16$, $p = 0.69$; social status: $\chi^2(2,21) = 4.61$, $p = 0.09$).

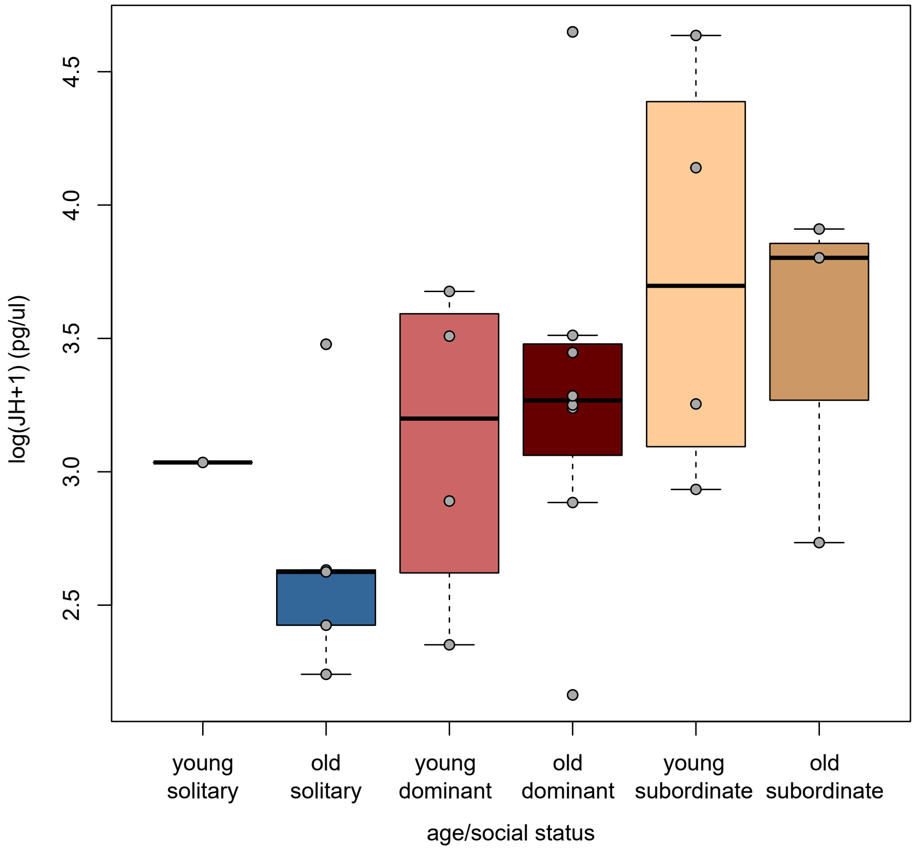


Fig. S18. Boxplot (median ± interquartile range) of log-transformed juvenile hormone titres (pg/µl) for females of different age and social status, including *E. dilemma* individuals. Points indicate individual titres. Juvenile hormone titres differed slightly according to social status when tested across all individuals, with solitary females exhibiting lower titres than females from social nests (LMM, age: $\chi^2(1,25) = 0.04$, $p = 0.84$; social status: $\chi^2(2,25) = 6.41$, $p = 0.04$).
